# HydroShoot: a functional-structural plant model for simulating hydraulic structure, gas and energy exchange dynamics of complex plant canopies under water deficit - application to grapevine (*Vitis vinifera* L.)

**DOI:** 10.1101/542803

**Authors:** R. Albasha, C. Fournier, C. Pradal, M. Chelle, J. A. Prieto, G. Louarn, T. Simonneau, E. Lebon

## Abstract

This paper aims at presenting HydroShoot, a functional-structural plant model (FSPM) that is developed to simulate gas-exchange rates of complex plant canopies under water deficit conditions, by scaling up gas-exchange rates from the leaf to the canopy levels. The main hypothesis is that simulating both the hydraulic structure of the shoot together with the energy budget of individual leaves is the asset for successfully achieving this up-scaling task. HydroShoot was hence built as the ensemble of three interacting modules: *hydraulic* which calculates the distribution of xylem water potential across shoot hydraulic segments, *energy* which calculates the complete energy budget of individual leaves, and *exchange* which calculates net assimilation and transpiration rates of individual leaves. HydroShoot was coupled with irradiance interception and soil water balance models, and was evaluated on virtual and real grapevines having strongly contrasted canopies, under well-watered and water-deficit conditions. HydroShoot captured accurately the impact of canopy architecture and the varying soil water deficit conditions on plant-scale gas-exchange rates and leaf-scale temperature and water potential distributions. Both shoot hydraulic structure and leaf energy budget simulations were, as postulated, required to adequately scaling-up leaf to canopy gas-exchange rates. Notwithstanding, simulating the hydraulic structure of the shoot was found far more necessary to adequately performing this scaling task than simulating leaf energy balance. That is, the intra-canopy variability of leaf water potential was a better predictor of the reduction of whole plant gas-exchange rates under water deficit than the intra-canopy variability of leaf temperature. We conclude therefore that simulating the shoot hydraulic structure is a prerequisite if FSPM’s are to be used to assess gas-exchange rates of complex plant canopies as those of grapevines. Finally HydroShoot is available through the OpenAlea platform (https://github.com/openalea/hydroshoot) as a set of reusable modules.

**Highlights:**

- Plant-scale gas-exchange rates were accurately reproduced using an FSPM based on the leaf-scale.
- Simulating the hydraulic structure of the shoot improves the prediction of plant gas-exchange rates under water deficit conditions.
- Simulating the intra-canopy variability of leaf temperature has a minor impact on plant-scale gas-exchange dynamics.
- Accurate predictions of gas-exchange rates of complex grapevine canopies require accounting for their hydraulic structure.

## Introduction

Climate change is seriously challenging viticulture sustainability in its current areas of production **(Hannah et al., 2013; van Leeuwen et al., 2013; Duchêne et al., 2014)**. One efficient short-term solution for hampering the projected adverse effects of water and heat stress on viticulture, is to reconsider training systems so that they allow maximizing the ratio of carbon assimilation (*A_n, plant_*) to water loss by transpiration (*E_plant_*) whilst maintaining optimal leaf temperature conditions **(Medrano et al., 2012; Duchêne et al., 2014; Palliotti et al., 2014)**. However, training systems present a wealth of possibilities **(Reynolds and Vanden Heuvel, 2009)** that cannot be compared experimentally. Therefore, the design of canopy structures that are adapted to adverse environmental conditions is most efficiently performed with the aid of models able to accurately predict the influence of canopy architecture on its gas exchange rates and leaves temperature under combined water and heat stress. That is what functional-structural plant models (FSPMs) offer **(Vos et al., 2010)**.

FSPMs received nevertheless little attention in grapevine scientific literature to assess the impact of shoot architecture on plant gas-exchange rates **(Medrano et al., 2015a)**. This is probably due to the inherent complexity in scaling up eco-physiological processes from the leaf to the canopy level, as strong variability in gas exchange rates (CO_2_ versus water vapor) exists inside the canopy driven by variations in micrometeorological conditions and leaf functional traits **(Niinemets et al., 2014; Medrano et al., 2015b)**. This scaling-up task is even more complex under water deficit conditions, as stomatal aperture is likely to reduce under water deficit in a non-uniform pattern across the canopy (e.g. **Gonzalez-Dugo et al., 2012; Ngao et al., 2017**) further distorting intra-canopy gas-exchange and leaf temperature variability **(Reynolds and Vanden Heuvel, 2009). Bauerle et al. (2007)** indicated that disregarding this variability may lead to strongly overestimate the predicted whole canopy daily transpiration flux, up to 25% greater than observed values as they found on a study on Red Maple (*Acer rubrum* L.)

Hence, adequately predicting the intra-canopy variability of both leaf stomatal conductance and temperature stands out as the key challenge to using FSPMs to predict plant gas-exchange rates under water deficit conditions. Yet, the remaining question is how can this variability be accurately described and what are their main determinants.

Describing the intra-canopy variability of both leaf stomatal conductance and temperature requires from the one hand to explicitly describe their drivers as a function of leaf position inside the canopy, and from the other hand to adequately account for their mutual interactions (how stomatal aperture affects leaf energy budget and *vice versa*) **(Chelle, 2005)**. The main drivers for both processes are commonly determined as incident shortwave irradiance, air temperature, air humidity and leaf “water status” which determines stomatal closure **(Damour et al., 2010)**. Among these drivers, leaf water status (which controls stomatal aperture and consequently both leaf transpiration and temperature) still makes no consensus in the scientific literature when it comes to determining what it refers to. A basal approach considers that leaf water status is equal to the soil water status (e.g. **Misson et al., 2004; van Wijk et al., 2000**), considering that the “remote” action of available water in the rhizosphere impacts uniformly all leaves, regardless of their position. By contrast, leaf water status may also be considered as a “local” water status specific to each individual leaf (e.g. **Tuzet et al., 2003; Buckley et al., 2003**) which results from the interplay between water demand (transpiration) and offer (xylem flow) at the leaf-scale, which are notably determined by the shoot hydraulic structure. This “local” approach mostly agrees with observations whereby stomatal closure is uneven across the canopy. Sunlit leaves, for instance, experience stronger water deficit than shaded leaves, and are therefore the first to undergo reductions in gas exchange rates under water deficit **(Escalona et al., 2003)**. This “local” approach seems hence as the most adequate in FSPM’s, however, the “remote” modelling approaches also proved satisfactory when individually considered. (e.g. **Dauzat et al., 2001; Bailey et al., 2016; Ngao et al., 2017**). It is hence unclear in literature how the simulation of leaf water status of individual leaves affects the predicted gas-exchange rates and leaf temperature distribution in FSPMs under water deficit and this matter needs to be assessed if FSPMs are to be used under water deficit conditions.

The few number of the existing grapevine FSPM’s in literature does not account for the interactions between water status, energy budget and gas-exchange rates at the leaf scale. **Prieto et al. (2012)** were probably the first to use an FSPM to examine the effects of canopy architecture on gas-exchange in grapevine (cv. Syrah). The authors coupled the grapevine-specific structural plant model proposed by **Louarn et al. (2008)** to leaf-level models of photosynthesis **(Farquhar et al., 1980)** and stomatal conductance (Leuning, 1995) but did not incorporate the effects of (soil) water deficit. More recently, Zhu et al. (2017) developed a model similar to that proposed by **Prieto et al. (2012)** including the effect of water deficit on gas-exchange and leaf temperature. Nevertheless, this model assumed a uniform xylem water potential across the shoot, that disregarded how shoot hydraulic structure affects leaf-scale gas-exchange rates, which is an unrealistic assumption for large plants when considering the substantial hydraulic resistances observed in the stems of mature grapevine **(Jacobsen et al., 2012)**. In addition, longwave energy exchange among leaves from the one hand, and between leaves and the surrounding elements from the other hand, were disregarded, which makes the application of this model to open field conditions not suitable since sky and soil longwave energies substantially affect leaves temperature **(Nobel, 2005)**. This has been solved in two models that link a complete energy budget with gas-exchange in perennials (**Bailey et al., 2016** for *Vitis vinifera* L. and *Acer x fremanii*; **Ngao et al., 2017** for *Malus pumila* Mill.) Yet again, both models were built at the leaf-cluster scale which does not allow accounting for the location of individual leaves in plant hydraulic structure necessary to calculate local leaf water status. In addition, both models consider the “remote” action of the rhizosphere on stomatal aperture instead of the “local’ water status, assuming again as negligible the potential contribution of shoot hydraulic structure on shaping the intra-canopy variability of leaf stomatal conductance and temperature.

In this paper, it is postulated that intra-canopy variability in both leaf water potential and leaf temperature are the main drivers for adequately predicting photosynthesis and transpiration fluxes at the plant scale under water deficit conditions using FSPMs. This paper has threefold objective. The first is to describe HydroShoot, a leaf-scale-based FSPM that allows predicting whole plant transpiration and photosynthesis rates by scaling-up these processes from the leaf-level. This model uses as lever for this scaling process the simulation of the interactions between hydraulic structure of the shoot, the energy balance, and gas-exchange rates of individual leaves. The second objective is to evaluate the performance of the model using both virtual and real canopies with data collected on photosynthesis and transpiration rates (plant scale), and stomatal conductance and temperature (leaf scale). Finally, the third objective is to examine how detailed hydraulic structure and energy budget simulations determine the predicted gas-exchange rates at the plant scale under water deficit conditions.

## Materials and methods

### Model structure and basic assumptions

HydroShoot is a static FSPM (with regards to plant structure) that takes plant shoot architecture, weather, and soil water conditions as inputs, and returns transpiration and net photosynthesis rates both of individual leaves and the whole plant at hourly time steps as outputs. It is conceived as a set of three modules which simulate water potential (*hydraulic* module), energy budget (*energy* module), and C_3_-type gas-exchange rates (*exchange* module). These three modules run jointly, having leaf xylem water potential and temperature at the leaf level as pivots (cf. *implementation and numerical solution* section). The formalisms used in each module are developed in the following sections.

### *Hydraulic* module

The *hydraulic* module computes water potentials of plant segments (output of the module) as a function of water flow in the plant and water potential of the soil (input to the module). The whole plant is compartmentalized in elementary conducting elements corresponding to petioles, internodes of the current-year stems, and elements of previous-year trunk and branches (internodes or pruning complexes). Leaves are treated in this system as nodes letting water flow but having no gradient in their water potential (*Ψ_leaf_*).

Water transfer across the hydraulic segments is simulated by analogy to Ohm’s law in electrical circuits (Figure 1). Each segment is characterized by its length (*L, m*) and hydraulic conductivity (*K, kg s^−1^ m MPa^−1^*), and is crossed by a water flux (*F, kg s*^−1^) which, together with conductivity modifies water head at its upper (downstream, *H_u_*) and lower (upstream, *H_l_*) extremities [*MPa*]:

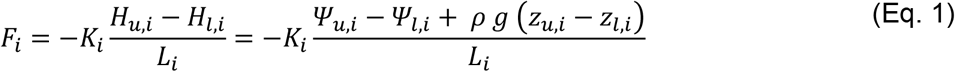

where *i* denotes the segment identifier, *Ψ_u, i_* and *Ψ_l, i_* are respectively xylem water potential at the upper and lower extremities, *z_u, i_* and *z_l, i_* are elevations of upper and lower extremities [*m*], respectively, *ρ* is water density [*kg m*^−3^] and *g* is the gravitational acceleration [*m s*^−2^].

**Figure 1:**
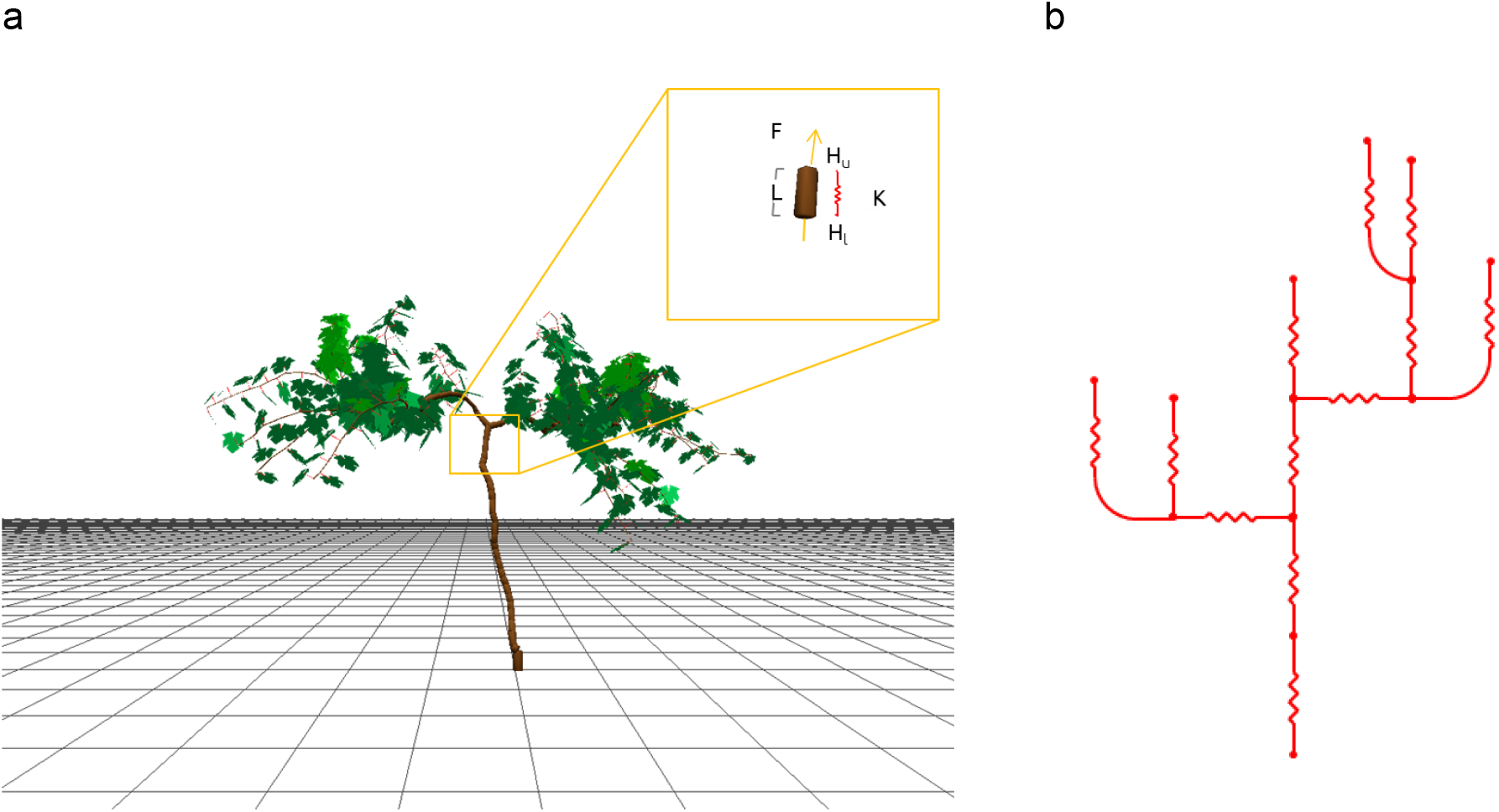
(a) Illustration of the parameters involved in hydraulic structure calculation of a stem segment: *F*, rate of water flow [*kg s^−1^*], *K*, hydraulic conductivity per unit stem length [*kg s^−1^ m MPa^−1^*], *L*, length of stem segment or internode [*m*] and *H_u_* and *H_l_* are respectively water potentials at upper (downstream) and lower (upstream) extremities of the stem segment [*MPa*]; and (b) the schematic representation of the electrical analog.

Xylem conductivity varies with water potential as a result of xylem cavitation under water deficit **(Tyree and Sperry, 1989)**. This relationship is described using a sigmoidal function:

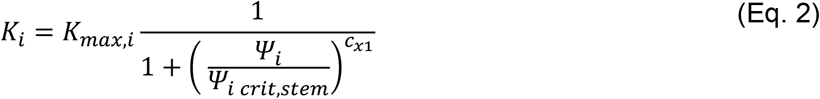

where *K_max, i_* is the maximum segment conductivity [*kg s*^−1^ *m MPa*^−1^], *Ψ_i_* is the arithmetic mean of water potential values of the segment 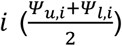 and *c*_*x*1_ [−] are shape parameters. *K_max, i_* is estimated empirically as proposed by **Tyree and Zimmermann (2002):**

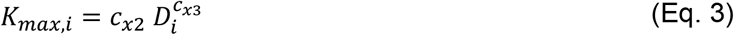

where *D_i_* is segment average diameter [*m*] and *c*_*x*2_ and *c*_*x*3_ are dimensionless shape parameters, mostly given within the ranges of [2.5, 2.8] and [2.0, 5.0], respectively **(Tyree and Zimmermann, 2002)**.

Equations 1 to 3 apply to all conducting segments (not leaves blades). Water potential of the upper extremity of the petiole is assumed equal to that of the lumped leaf water potential *Ψ_leaf_*.

### *Exchange* module

The *exchange* module computes the rates of net carbon assimilation and transpiration per unit surface area (respectively *A_n_* and *E*) for each individual leaf as a function of micrometeorological conditions and leaf water status. The calculations use the analytical solution proposed by **Yin et al. (2009)** for coupling the *C_3_* photosynthesis model of **Farquhar et al. (1980)** to the stomatal conductance model of **Ball et al. (1987)**. This coupling is based on Fick’s first law of diffusion, whereby *A_n_*, the stomatal conductance to CO_2_ (*g_s, co_2__*), and the mesophyll conductance (*g_m_*) are used. However, as Farquhar’s model has been thoroughly detailed in literature, its description is given in Appendix I. The focus of this section is given instead to the stomatal conductance formulae which are a key element in this work.

*g*_*s, CO*_2__ is calculated according to **Yin et al. (2009)** as:

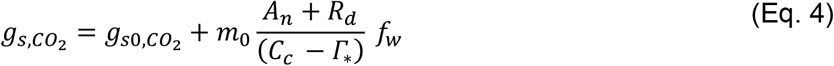

where *g*_*s*0, *CO*_2__ is the residual stomatal conductance to *CO_2_* [*mol*_*CO*_2__ *m*^−2^ *s*^−1^], *R_d_* is mitochondrial respiration in the light [*μmol*_*CO*_2__ *m*^−2^ *s*^−1^], *Γ*_*_ is *CO*_2_ compensation point in the absence of mitochondrial respiration [*μmol*_*CO*_2__ *mol*_*CO*_2__ ^−1^], *m*_0_ is a dimensionless shape parameter, and *f_w_* is a dimensionless function representing the response of *g*_*s, CO*2_ to air water vapor deficit (*VPD, kPa*). *f_w_* is deduced from the stomatal conductance model of Leuning (1995) as:

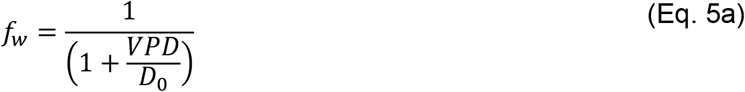

where *D*_0_ is a scaling parameter [*kPa*].

Eq. 5a does not account for stomatal sensitivity to soil water deficit (“remote” approach) or local leaf water potential (“local” approach). **Tuzet et al. (2003)** and **Leuning et al. (2004)** suggested to express *f_w_* as a function of the local *Ψ_leaf_*. This function is implemented in HydroShoot following **Nikolov (1995):**

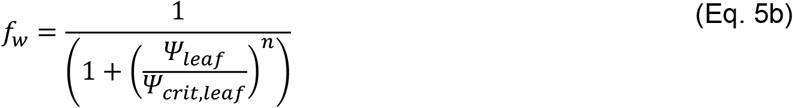

where *Ψ_crit, leaf_* is a critical leaf water potential threshold [*MPa*] at which stomatal conductance is reduced by 50%, and *n* is a shape parameter [–]. The same last equation is used to express the dependency of *g*_*s*0, *CO*_2__ on the remote soil water potential (*Ψ_soil_*):

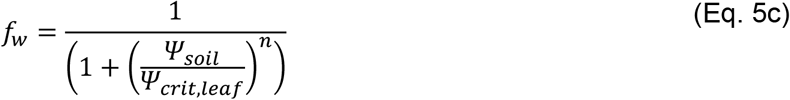

The transpiration rate *E* [*mol*_*H*_2_*O*_ *m*^−2^ *s*^−1^] is calculated as:

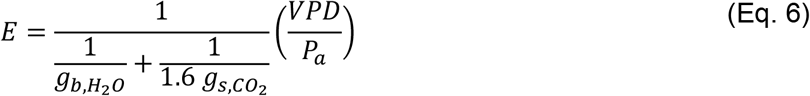

where *P_a_* is the atmospheric pressure [*MPa*] and *g*_*b, H*_2_O_ is the boundary layer conductance to water vapor [*mol*_*H*_2_O_ *m*^−2^ *s*^−1^], described by **Nobel (2005)** as:

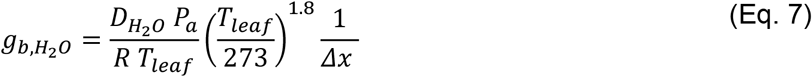

where *D*_*H*_2_O_ is the diffusion coefficient of H_2_O in the air at 0 *°C* (2.13*10^−5^ *m*^2^ *s* ^−1^), *P*_0_ is the ambient air pressure at 0 *°C* temperature [*MPa*], and *Δx* is the thickness of the boundary layer [*m*] which is defined as **(Nobel 2005)**:

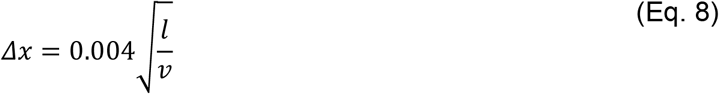

where *l* is the mean length of the leaf in the downwind direction [*m*], set to 70% of blade length, and *v* is the ambient wind speed [*m s*^−1^].

Finally, mesophyll conductance to CO_2_ is assumed to simply depend on bulk leaf temperature **(Evers et al., 2010)** following an Arrhenius equation trend (as for photosynthetic parameters, cf. Eq. A8) with a basal value at 25 *°C* set to 0.1025 [*mol*_*CO_2_*_ *m*^−2^ *s*^−1^].

#### Intra-canopy variability in photosynthetic capacity

Leaf photosynthetic traits (maximum carboxylation rate *V_c max_*, maximum electron transport rate *J_max_*, triose-phosphate transport rate *TPU* and *R_d_*; cf. Appendix I) have been shown to strongly vary within the plant canopy as a result of the variability in micrometeorological conditions **(Niinemets et al., 2014)**. HydroShoot accounts for this variability by considering leaf nitrogen content per unit leaf surface area (*N_a_, g_N_ m^−2^*) as the pivotal trait to determine the photosynthetic capacity of leaves **(Prieto et al., 2012)** as follows:

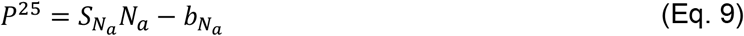

where *P^25^* is the value at 25 *°C* for any of the rates *V_c max_, J_max_*, *TPU* and *R_d_* (given as inputs), and *S_N_a__* [*μmol_CO_2__ g_N_^−1^ s^−1^*] and *b_N_a__* [*μmol_CO_2__ m^−2^ s^−1^*] are the slope and the intercept of the linear relationship with *N_a_* specific to each rate. *N_a_* is calculated as the product of nitrogen content per unit leaf dry mass *N_m_* [*g*_*N*_ *g*_*dry matter*_^−1^] and leaf dry mass per area *LMA* [*g_dry matter_ m^−2^*]. *N_m_* linearly varies with plant age, expressed as the thermal time cumulated from budburst (input of the model), and *LMA* is determined by leaf exposure to light during the last past days **(Prieto et al., 2012)**. This is expressed respectively in the two following equations:

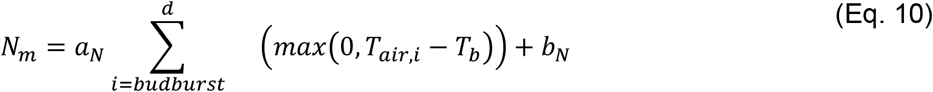

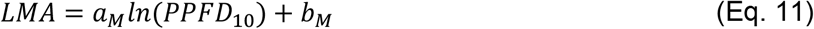

where *T_air, i_* is the mean temperature of the day *i* [°*C*] and *T_b_* is the base temperature (minimum required for growth) [°*C*], set to 10°C for grapevine and used for the calculation of thermal time since budburst, *a*_*N*_[*g*_*N*_ *g*_*dry matter*_^−1^ °*Cd*^−1^] and *b_N_* [*g_N_ g_dry matter_^−1^*] are the slope and intercept of the linear relationship between *N_m_* and accumulated thermal time since budburst, *PPFD*_10_ [*mol*_*photon*_ *m*^−2^ *d*^−1^] is the cumulative photosynthetic photon flux density irradiance intercepted by the leaf (output of the energy module) averaged over the past 10 days, *a_M_* [*g_dry matter_ m*^−2^] and *b_M_* [*g_dry matter_ m*^−2^] are the slope and intercept of the linear relationship between *LMA* and the logarithm of *PPFD_10_*.

Finally, this module was provided with photoinhibition model as this phenomenon is frequently reported to affect grapevines under combined heat and water stresses **(Correia et al., 1990; Flexas and Medrano, 2002; Lovisolo et al., 2010)**. The simple photoinhibition model implemented in HydroShoot is detailed in Appendix II and assumes that combined heat and water stresses inhibit photosynthesis by reducing the electron transport rate (cf. *J* in Eq. A6) as the result of an increased of deactivation energy *ΔH_d_* (cf. equations A9 and A10).

### *Energy* module

The *energy* module computes the temperature of individual leaves based on a detailed energy balance model that is developed in the supplementary materials provided with this paper and only briefly described hereafter for the sake of simplicity.

Each leaf is represented as a group of solid flat triangles. It gains energy from the absorbed shortwave (solar irradiance) and thermal longwave irradiance from the sky, the soil, and the neighbouring leaves (indexed *j*). It loses energy through its own emission in the thermal longwave band and through latent heat due to transpiration (output of *exchange* module). Finally, it exchanges energy with the surrounding air by thermal conduction-convection. The resulting leaf-scale energy balance equation writes:

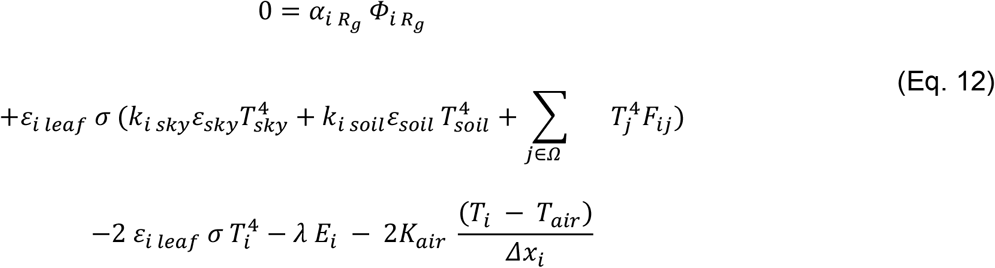

where *i* refers to leaf identifier, *α_Rg_* is lumped leaf absorptance in the shortwave band [–], *Φ_Rg_* is flux density of shortwave global irradiance *R_g_* [*W m*^−2^], *∊_leaf_*, *∊_sky_* and *∊_soil_* are emissivity-absorptivity coefficients of the leaf, sky and soil, respectively [–], *σ* is the Stefan-Boltzmann constant [*W m*^−2^ *K*^−4^], *T_Sky_*, *T_soil_*, and *T_air_* are respectively the sky, soil, and air absolute temperatures [*K*] all taken as input parameters for HydroShoot; *T_j_* is temperature of neighboring leaf *j* (solved by convergence, see Implementation and numerical resolution section), *λ* is latent heat for vaporization [*W s mol*^−1^], *K_air_* is the thermal conductivity of air [*W m*^−1^ *K^−1^*], and finally, *k_sky_, k_soil_* and *F_ij_* are the form factors of the sky, soil and canopy elements in the sphere *Ω* surrounding the leaf *i* **(Chelle et al., 1998)**. *α_Rg_*, *ρ_TIR_* and *∊_leaf_* are input parameters considered as uniform for all leaves.

Since the resolution of the last equation is highly time-consuming, we assumed the energy gain from the neighbouring leaves through thermal longwave as a lumped term whereby average leaf temperature *T_leaves_* is considered instead of individual leaves **(Dauzat et al., 2001)**. In this case, the lumped form factor *Σ_j∊Ω_ F^ij^* is simply taken as 1 – (*k_sky_* + *k_soil_*) (that is the solid angle where neither the sky nor the soil are seen by a single leaf). The former equation becomes:

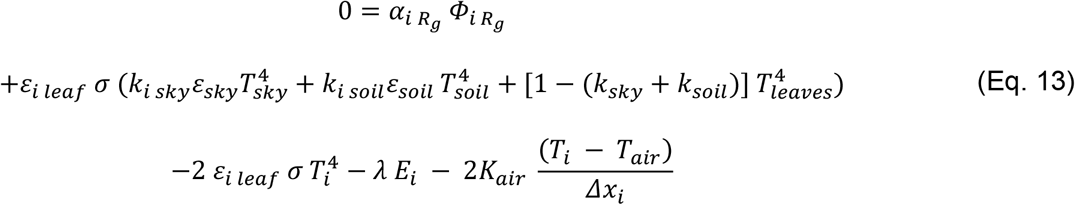

### Implementation and numerical resolution

HydroShoot is developed using the *Python* programming language (Python Software Foundation http://www.python.org) in the OpenAlea platform **(Pradal et al., 2008; Pradal et al., 2015)**. It uses the Multiscale Tree Graph (MTG) method **(Godin and Caraglio, 1998; Balduzzi et al., 2017)** as a central data-structure in order to allow indirect communication between the different models which favor modularity **(Fournier et al., 2010; Garin et al., 2014)**. Each process has been implemented as a reusable component in OpenAlea and can be reused independently in other models and composed in various ways, provided that the other models are written in the *Python* language.

The resolution of HydroShoot equations is performed by an iterative procedure that is schematized in Figure 2.

**Figure 2:**
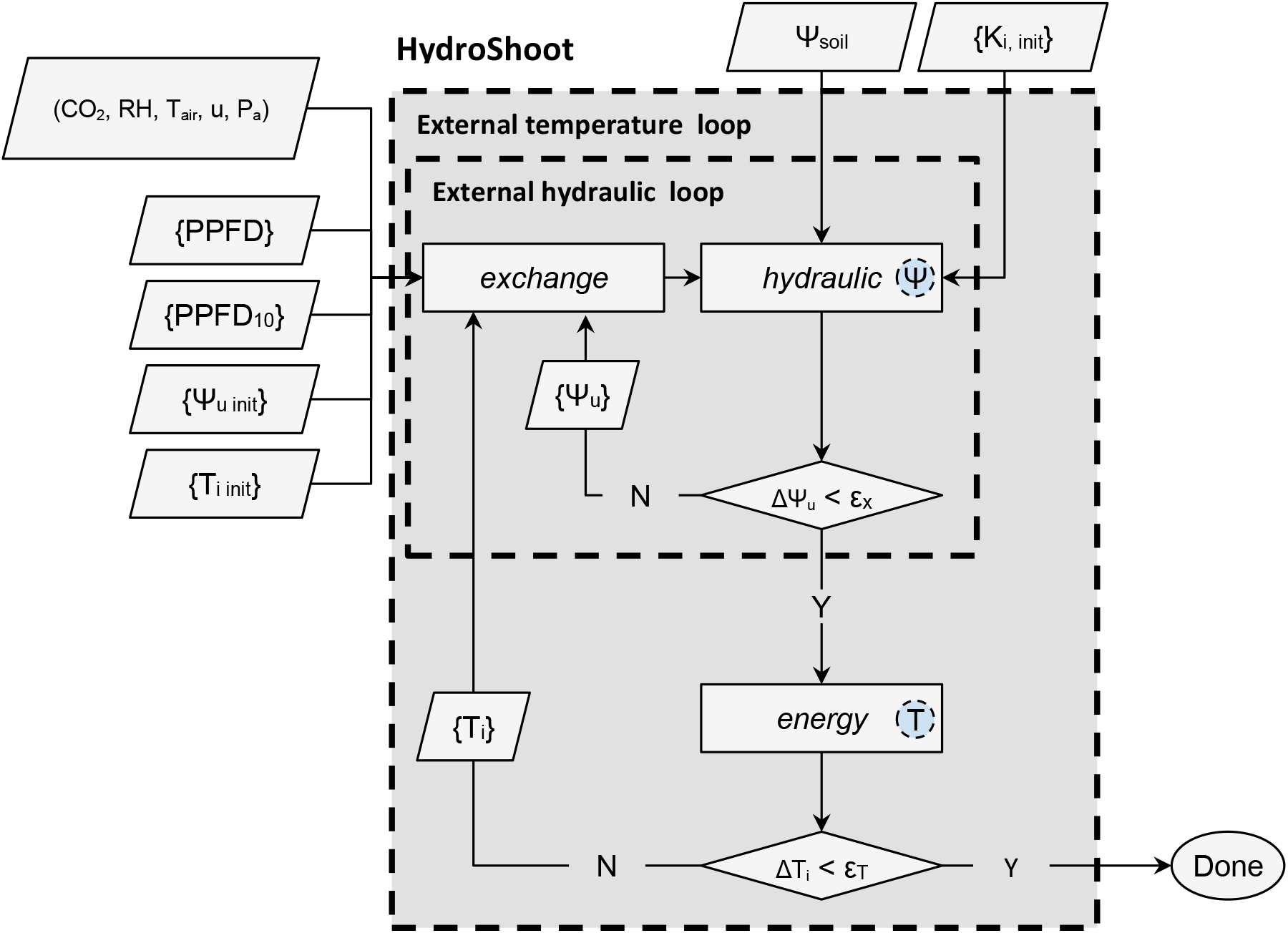
Schematic representation of the numerical resolution of HydroShoot. Meteorological inputs that are common to all leaves are air temperature (*T_air_, K*), air relative humidity (*RH*, %), air CO_2_ concentration [*μmol mol*^−1^], wind speed (*u, m s*^−1^), and atmospheric pressure (*Pa, kPa*). Inputs per individual leaves are the absorbed photosynthetic photon flux density (*PPFD*, *μmol m*^−2^ *s*^−1^) and *PPFD*_10_ the absorbed *PPFD* during the last 10 days. *Ψ_u_* is xylem water potential at the nodes between each pair of stem elements [*MPa*]. *Ψ_u init_* is initial *Ψ_u_* [*MPa*]. *Ψ_soil_* is soil water potential. *Ψ_u_* [*MPa*]. *T_i_* is leaf temperature [*K*]. *T_i init_* is initial *T_i_* [*K*]. *K_init_*[*k_g_ s*^−1^ *m MPa*^−1^] is initial hydraulic conductivity of each segment, *∊_x_* is the maximum allowable error of the estimation of xylem water potential [*MPa*] and *∊_T_* is the maximum allowable error of the estimation of leaf temperature [*K*]. Circles inside module boxes indicate internal iteration loops. Symboles between curly brackets represent spatially-structured variables.

Iterations have three levels. The first is in the *hydraulic* module and concerns calculating xylem water potential of plant segments in interaction with their hydraulic conductivity (interdependent processes, cf. equations 1 and 2). The second level is between the *exchange* and *hydraulic* modules in order to calculate jointly gas-exchanges rates and leaf water potential values (transpiration affects the hydraulic structure Eq. 1 while the latter affects stomatal aperture Eq. 5b). The third level is between the *energy* module and both *exchange* and *hydraulic* modules, so that at each time new transpiration fluxes are calculated, leaf temperature values are updated and the new temperature values are used to update gas-exchange rates which in their turn impose new xylem water potential distribution. The details on the numerical resolution are given in Appendix III.

### Coupling with irradiance and soil models

HydroShoot needs irradiance absorption by individual leaves and soil water potential as inputs. It is therefore coupled in this work to Caribu irradiance model **(Chelle et al., 1998)** and to a simple soil water-budget model in order to calculate respectively irradiance absorption (*PPFD*) and soil water potential (*Ψ_soil_*) values on an hourly basis (Figure 3).

**Figure 3:**
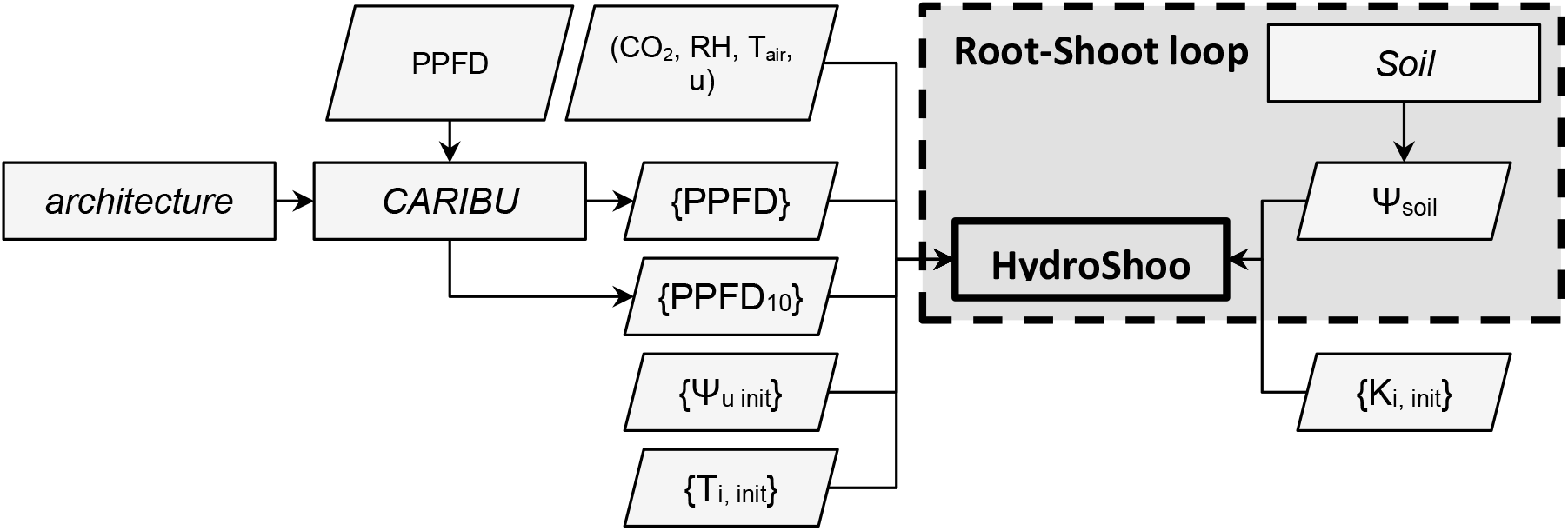
Flowchart of the modelling frame used in this application example. Meteorological inputs that are common to all leaves are air temperature (*T_air_, K*), air relative humidity (*RH*, %), air CO_2_ concentration [*μmol mol*^−1^], wind speed (*u, m s*^−1^), and atmospheric pressure (*Pa, kPa*). Inputs per individual leaves are the absorbed photosynthetic photon flux density (*PPFD*, *μmol m*^−2^ *s*^−1^) and *PPFD*_10_ the absorbed *PPFD* during the last 10 days. *Ψ_u, init_* is the initial xylem water potential at the nodes between each pair of stem elements [*MPa*]. *Ψ_soil_* is soil water potential. *T_i, init_* is initial temperature of individual leaves [*K*]. *K_init_*[*k_g_ s*^−1^ *m MPa*^−1^] is initial hydraulic conductivity of each segment. Symbols between curly brackets represent spatially-structured variables. *architecture, CARIBU* **(Chelle et al., 1998)** and *soil* are external modules used to simulate canopy architecture, irradiance interception, and soil water potential, respectively.

The *soil* module links transpired water rates to the transpirable soil water volume (*TSW*) in order to predict the hourly variations in *Ψ_soil_*. At the beginning of each calculation step, transpired water volume from the previous step is withdrawn from the *TSW*. The soil volumetric water content *Θ_soil_* is then determined by dividing *TSW* by the effective soil porosity. *Ψ_soil_* is then obtained from *Θ_soil_* from the water retention curve **(van Genuchten, 1980)** and used as an input for the *hydraulic* module. This procedure is referred to as the *Root-Shoot loop* in Figure 3.

## Model evaluation

Model evaluation was performed in three steps. Firstly, the coherence between expected and simulated gas-exchange, temperature, and xylem water potential dynamics for different canopies architectures was assessed. Secondly, the precision was assessed by comparing model outputs to measured plant gas-exchange rates and leaf stomatal conductance, water potential and temperature. Finally, the required complexity level was evaluated, whereby we sought at determining whether simulating the hydraulic structure and energy balance were (both) required in order to obtain accurate predictions of gas-exchange rates at the plant scale. For all the following simulations, parameter values are given in Appendix IV.

### Coherence

For this aim, HydroShoot was run on 3 virtual grapevine canopies which share the same soil type, soil initial water content, weather conditions, and total leaf area, and differ only in their shoot architecture (Figure 4).

**Figure 4:**
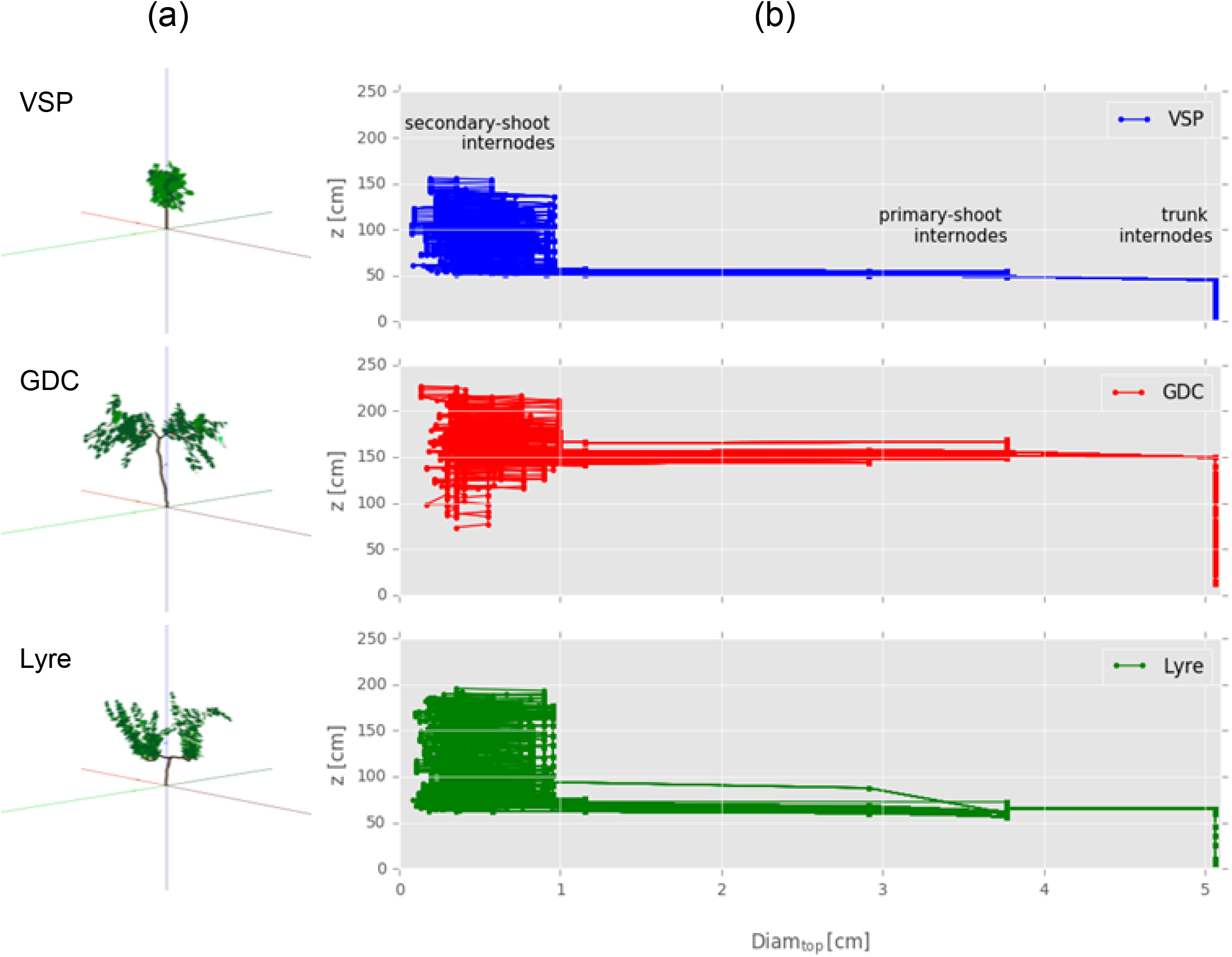
Mock-ups of three virtual grapevine canopies trained to Vertical Shoot Positioning (VSP), Geneva Double Curtain (GDC) and Lyre systems. Canopies are shown on the left column while conducting segments (primary and secondary internodes, petioles) diameters distribution is shown on the right column (Diamtop denotes the diameter of internodes).

The 3 canopies were trained on three different training systems: Vertical Shoot Positioning (VSP), Geneva Double Curtain (GDC) and Lyre systems (Figure 4a). All canopies had the same leaf area (5.7 *m^2^*), internode diameter distribution (Figure 4b), planting density (inter-and intra-row spacing of 3.6 and 1.0 *m* respectively), soil type (Sand Loam) and initial collar water potential (−0.6 *MPa*).

The simulations were run using weather data extracted from the database of the weather station of the National Institute for Agricultural Research (INRA) in Montpellier (3°53” E, 43°37” N, 44 m alt) on July 29^th^ 2009 (DOY 210). Weather conditions corresponded to a warm day having minimum and maximum air temperature of 19 and 34 °C respectively, relative humidity oscillating between 32 and 44%, wind speed at 2 *m* height going from 0 to 2 *m s^−1^*, and a clear sky with a maximum *PPFD* of 1670 *μmol m^−2^ s^−1^* at solar midday.

#### Precision

The precision of simulated outputs was evaluated by running HydroShoot on real canopies using collected data from experiments conducted in 2009 and 2012 on grapevine (cv. Syrah, grafted on SO4) at INRA, in Montpellier (same above-mentioned station). Five grapevines trained with two contrasting training systems were considered (cf. Figure 5): GDC in 2009 and VSP in 2012. Grapevine rows were oriented 140° from North on a shallow sandy loam soil with a low water holding capacity. Inter-row spacing was 3.6 *m* for GDC and 1.8 *m* for VSP. Intra-row spacing was 1 *m* (cf. supplementary materials for a detailed description of measurements).

**Figure 5:**
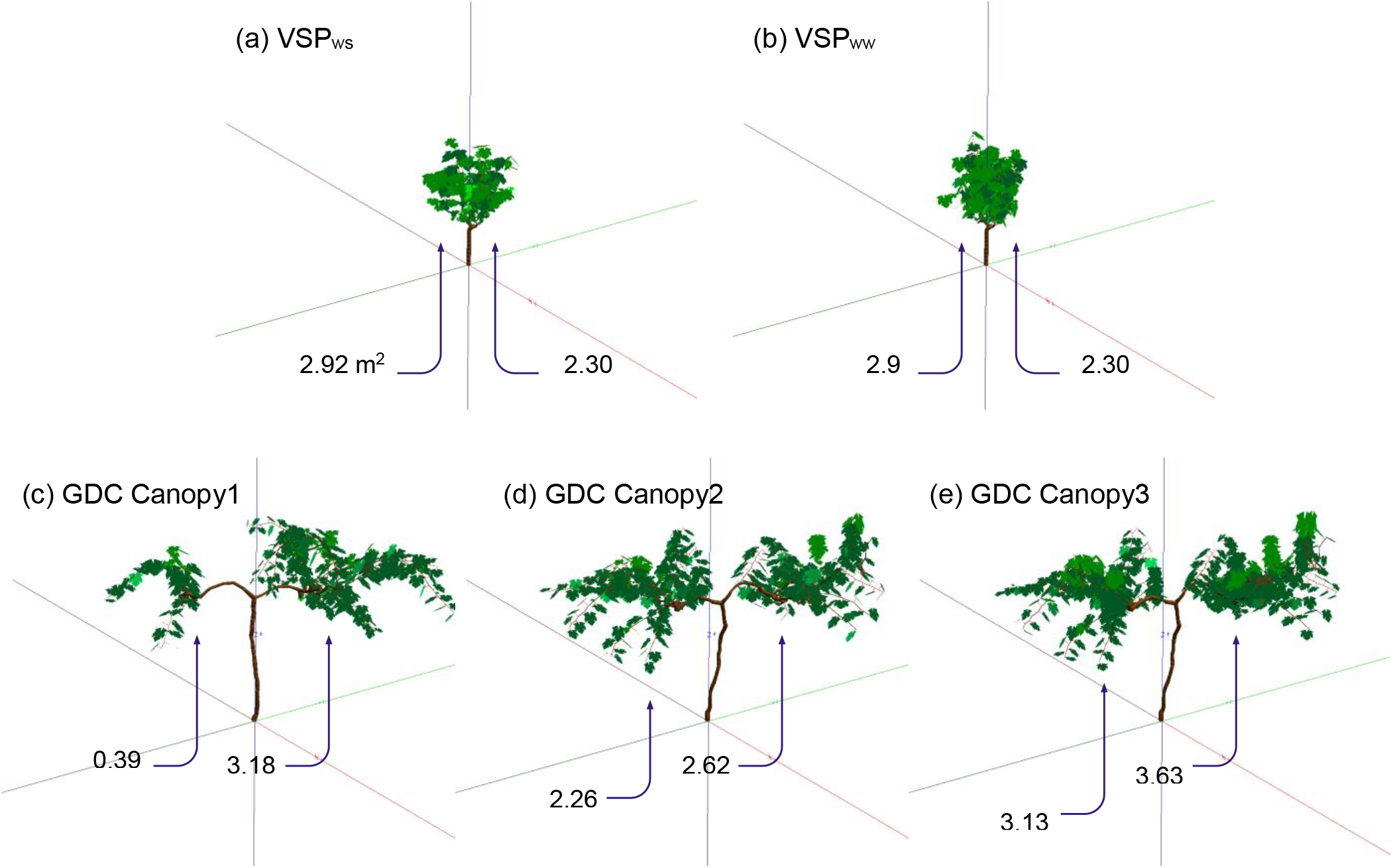
3D mockups of grapevines plants trained to Vertical Shoot Positioning system (VSP, a and b) under water deficit (VSP_ws_), well water conditions (VSP_ww_) and Geneva Double Curtain (GDC, c, d and e). The mockups were reconstructed from measured leaf surface profiles using the *architecture* module (input to HydroShoot). Numbers below canopies indicate leaf area per cordon [*m*^2^].

Data on VSP grapevines (2009) were collected during a period of 4 days under well-watered and water-deficit conditions. Water deficit was created by cutting off the irrigation system on the first day of the experiment (July 29^th^). Whole plant transpiration *E_plant_* and net assimilation *A_n, plant_* were monitored using open portable gas-exchange chambers **(Perez Peña and Tarara, 2004)**. Temperature of individual leaves were monitored using thermocouples inserted into the primary veins of 10 fully-developed individual leaves positioned on different heights from the top of the canopy to the inside, so that temperature gradient resulting from different irradiance conditions was captured.

Data on GDC grapevines (2012) were also collected during a 4 days experiment (starting on August 1^st^), but only under water-deficit conditions. Only *E_plant_* rate was monitored by measurements of sap flow installed on the two cordons of the GDC plants. Stomatal conductance and leaf water potential measurements were performed for a number of leaves on GDC grapevines during the experiment but the exact position of leaves was not reported with measurements.

For both VSP and GDC grapevines, shoot architecture was constructed based on digitisation data, using a grapevine-specific shoot architecture module following a Multiscale Tree Graph (MTG) approach **(Godin and Caraglio, 1998; Pradal et al., 2008; Balduzzi et al., 2017)**, in which organs topological connections and geometry were associated to shoot architecture (Figure 5). Plant mockups were produced so that the simulated vertical and horizontal profiles of leaf surface area fitted those observed.

### Complexity

In order to explore the contribution of HydroShoot’s *hydraulic* and *energy* modules components to the final simulation output, a sensitivity analysis was performed by plugging/unplugging each of these components and observing the resulting difference on simulated outputs. This procedure aims *in fine* at evaluating whether adding complexity to an FSPM would improve its performance in predicting gas-exchange dynamics at the plant-scale. The following simulation combinations are used:

- sim0: the reference (complete) version of HydroShoot having the ensemble of its components;
- sim1: stomatal conductance varies with *VPD* (as described by **Leuning, 1995** in Eq. 5a) regardless of leaf water potential;
- sim2: the hydraulic structure is disregarded (water potential of all leaves is forced equal to water potential at the collar) and stomatal conductance varies with collar water potential Eq. 5c;
- sim3: energy balance is disregarded, that is all leaves have the same temperature as that of the air;
- sim4: the same case of sim1 but using tighter control of *VPD* on stomatal conductance (*D*_0_ in Eq. 5a is set to 1 instead of 30 as proposed by **Prieto et al. (2012)**.

### Evaluation criteria

The overall adequacy between observed and simulated variables was assessed based on the estimation of the mean bias error (*MBE*) and root mean square error (*RMSE*):

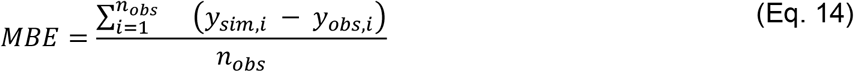

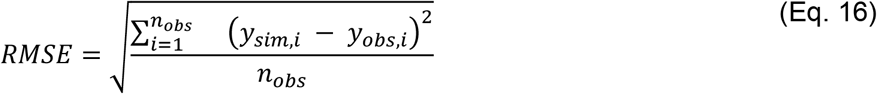

where *y_sim_* and *y_obs_* are respectively simulated and observed variables values and *n_obs_* is the number of observations.

## Results and discussion

### Outputs coherence

Simulation outputs for virtual canopies are illustrated in Figure 6 (plant-scale outputs) and Figures 7 and 8 (leaf-scale outputs).

**Figure 6:**
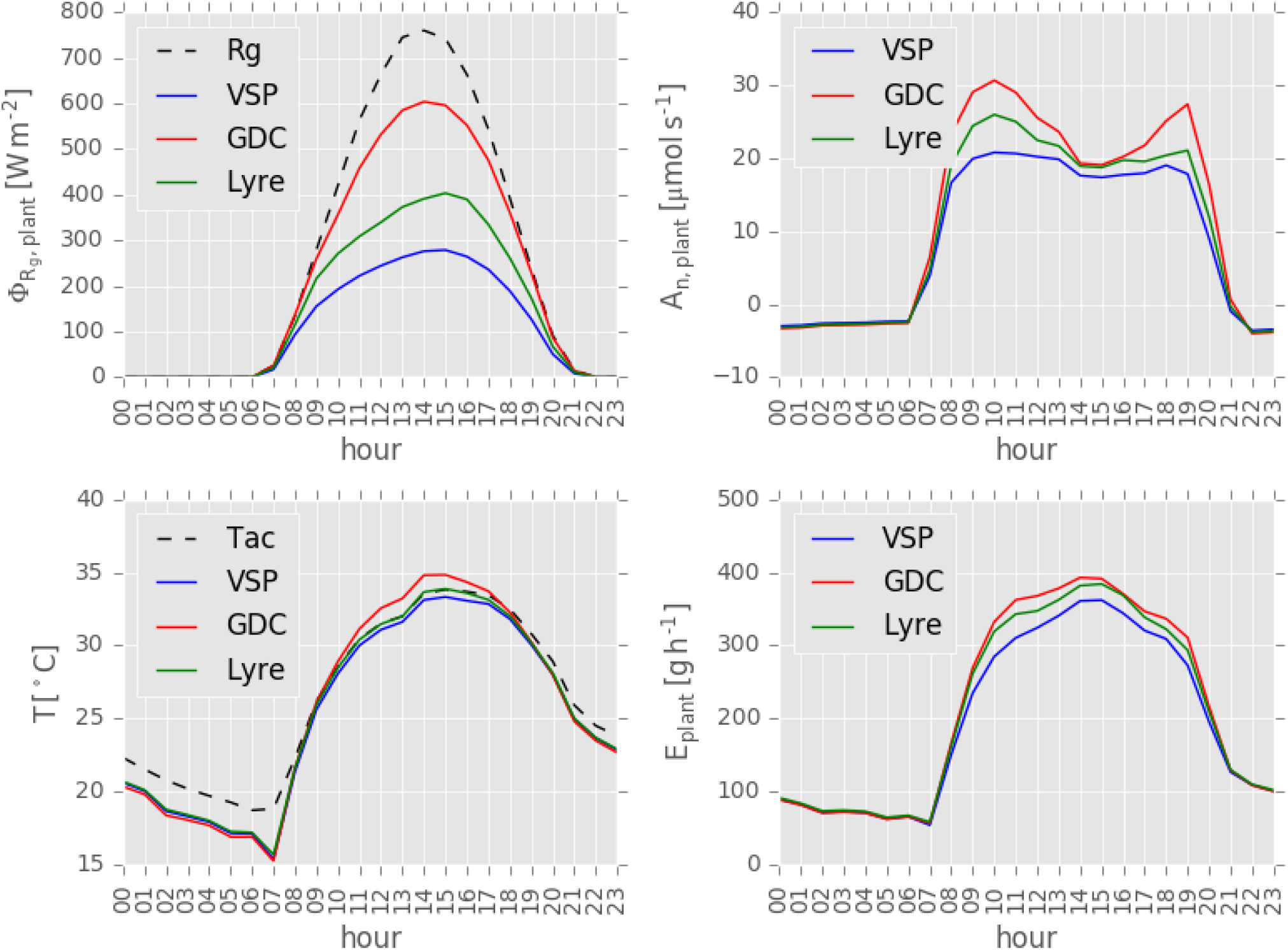
Simulation of absorbed irradiance (a), net carbon assimilation (b), temperature (c) and transpiration (d) at the plant-scale for three contrasted grapevine canopies (VSP, GDC and Lyre); *R_g_* is the incident global irradiance and *T_ac_* is air temperature. Temperature curves (c) trace the hourly values of the median of leaves temperatures.

The simulations at the plant scale (Figure 6) show that GDC canopies had the highest absorbed irradiance rates *Φ_Rg, plant_*, followed by Lyre and VSP canopies (Figure 6a), reflecting the higher exposure to solar irradiance using the GDC system. This trend was reflected on carbon assimilation *A_n, plant_* (Figure 6b), temperature *T* (Figure 6c), and transpiration *E_plant_* (Figure 6d), whereby highest values were obtained for GDC then Lyre followed by VSP canopies.

Midday depression in *A_n, plant_* was simulated for the three canopies proportionally to the absorbed *Φ_Rg, plant_*, that is highest for GDC and lowest for VSP (Figure 6b). The higher transpiration rates of GDC led to simulate lower leaf water potential values (Figure 7) around midday, which, combined with higher absorbed irradiance, led also to higher leaf temperatures (Figure 8). The combined effects of lower leaf water potential and higher temperature in GDC led to simulate a higher effect of midday depression in *A_n, plant_* compared to Lyre and VSP canopies as may be expected.

**Figure 7:**
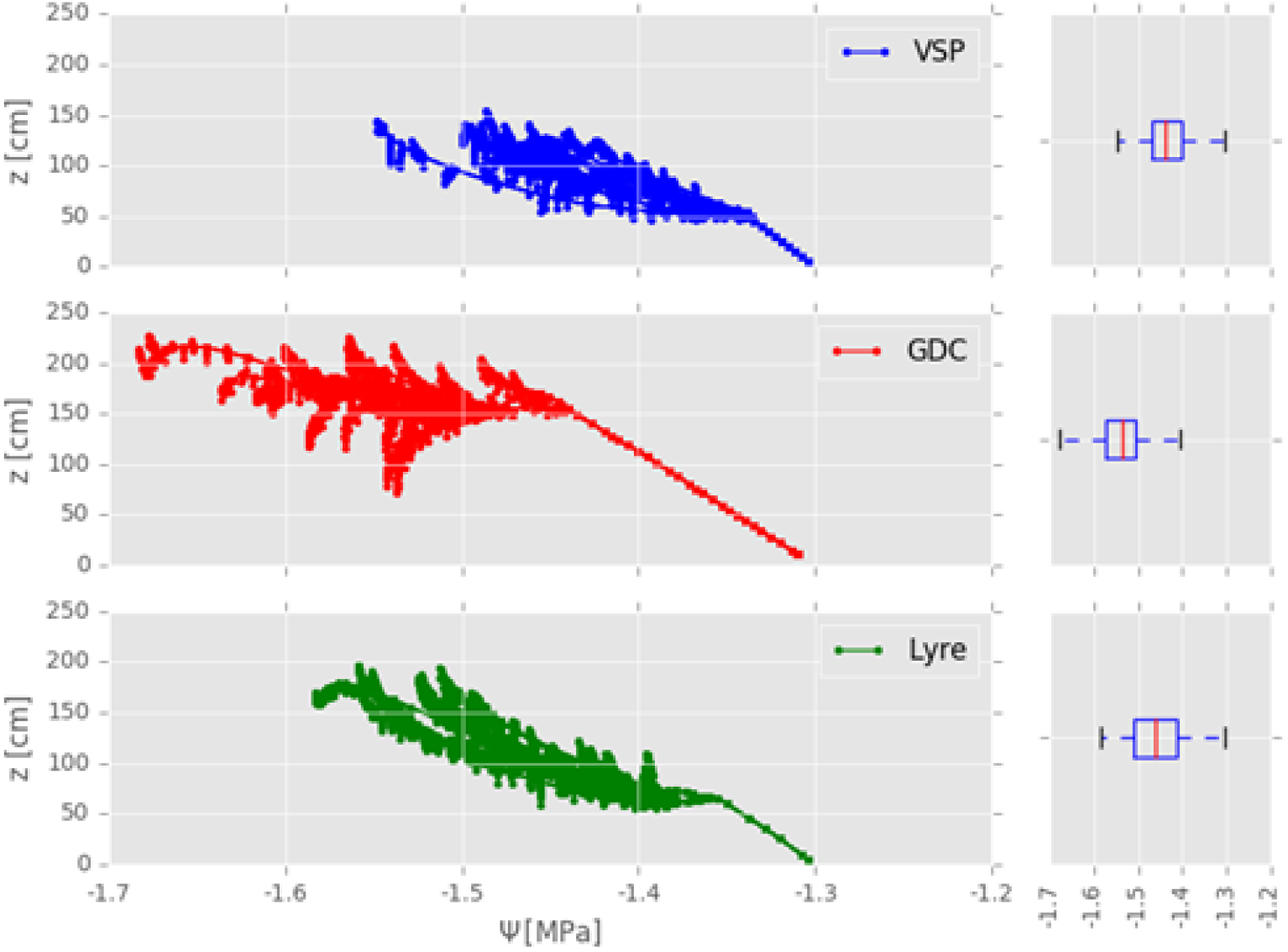
Snapshot at solar midday (14:00hs) of water potential distribution across the shoot (left column) and only for leaves (boxplots, right column) for three contrasted grapevine canopies (VSP, GDC and Lyre).

**Figure 8:**
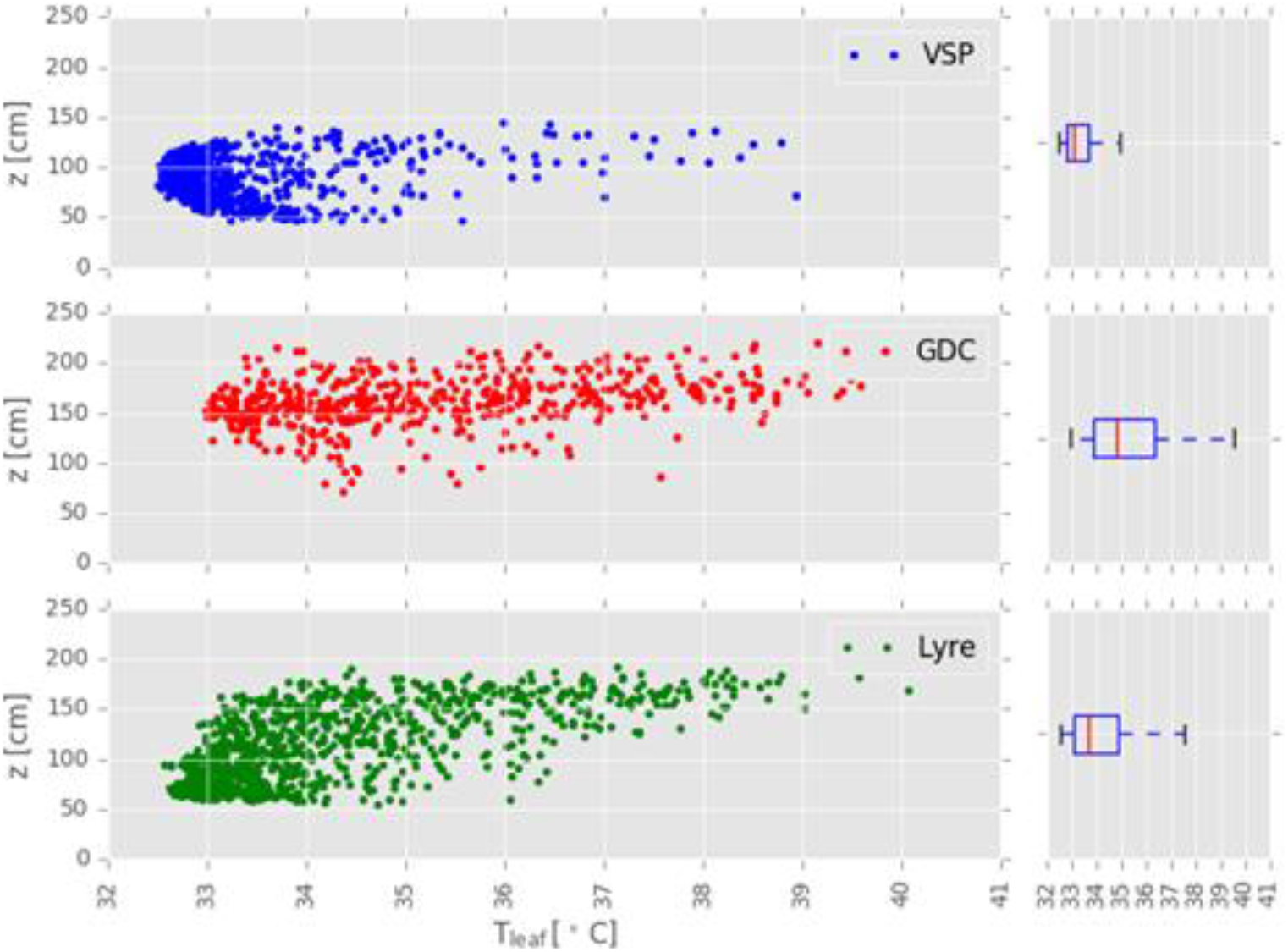
Snapshot at solar midday of individual leaf temperature values (left column) and leaf temperature distribution (boxplots, right column) for three contrasted grapevine canopies (VSP, GDC and Lyre).

This first illustrative example on virtual canopies shows that the effect of canopy architecture on its gas-exchange and temperature behaviour is captured in HydroShoot. The comparison to measurements in the following section will show how the observed dynamics on both plant and leaf scales are accurately reproduced using HydroShoot for two real canopies.

## Comparison to observed data

### Leaf-scale

Simulation results at the leaf-scale are shown in Figures 9, 10 and 11, respectively for water potential, stomatal conductance, and temperature. The dynamics of these variables were adequately reproduced but with some discrepancies regarding the onset timing of the effect of water-deficit. Due to a strong uncertainty on the exact position of leaves where measurements were performed, the comparison between observed and simulated processes at the leaf-scale will focus on variable dynamics rather than absolute values.

**Figure 9:**
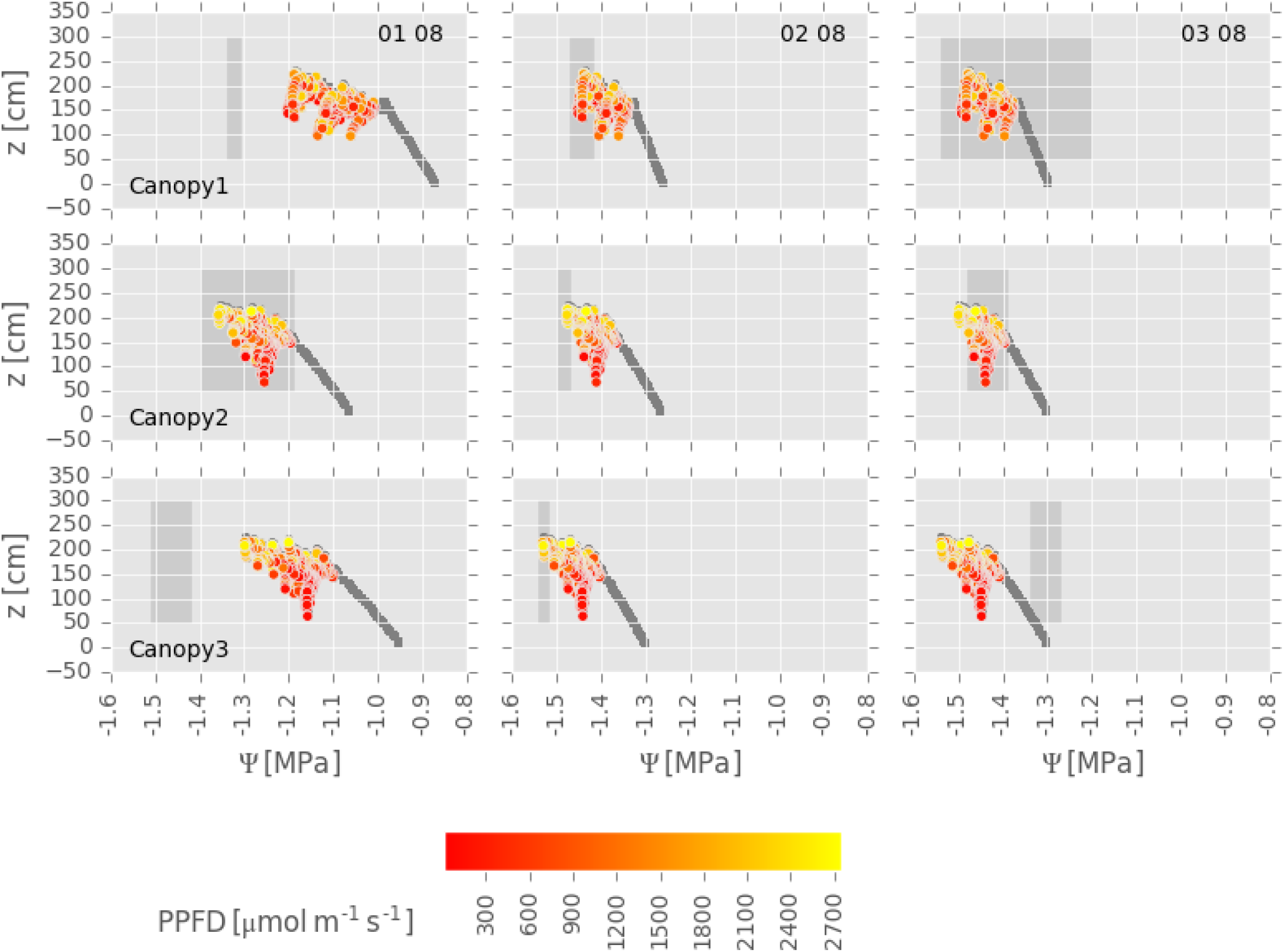
Snapshots of the simulated hydraulic structure of the three GDC canopies considered in this study, referred to as ‘Canopy1’, ‘Canopy2’ and ‘Canopy3’, respectively, prior to solar noon, during the first three days following the onset of soil water deficit, respectively 01 08, 02 08 and 03 08. Soil predawn water potential of the three days was equal to −0.19, −0.38, −0.61 *MPa* respectively. Filled circles represent xylem water potential, their colors (only for leaves) represent the absorbed *PPFD* value per unit leaf surface area; sunlit leaves are yellow while shaded leaves are red, and grey circles are for the trunk. Due to uncertainties in measurements locations, the observed water potential values of sunlit leaves are indicated by the grey patches which cover minimum and maximum leaf water potential values. Observed data were collected from experiments conducted in 2012.

**Figure 10:**
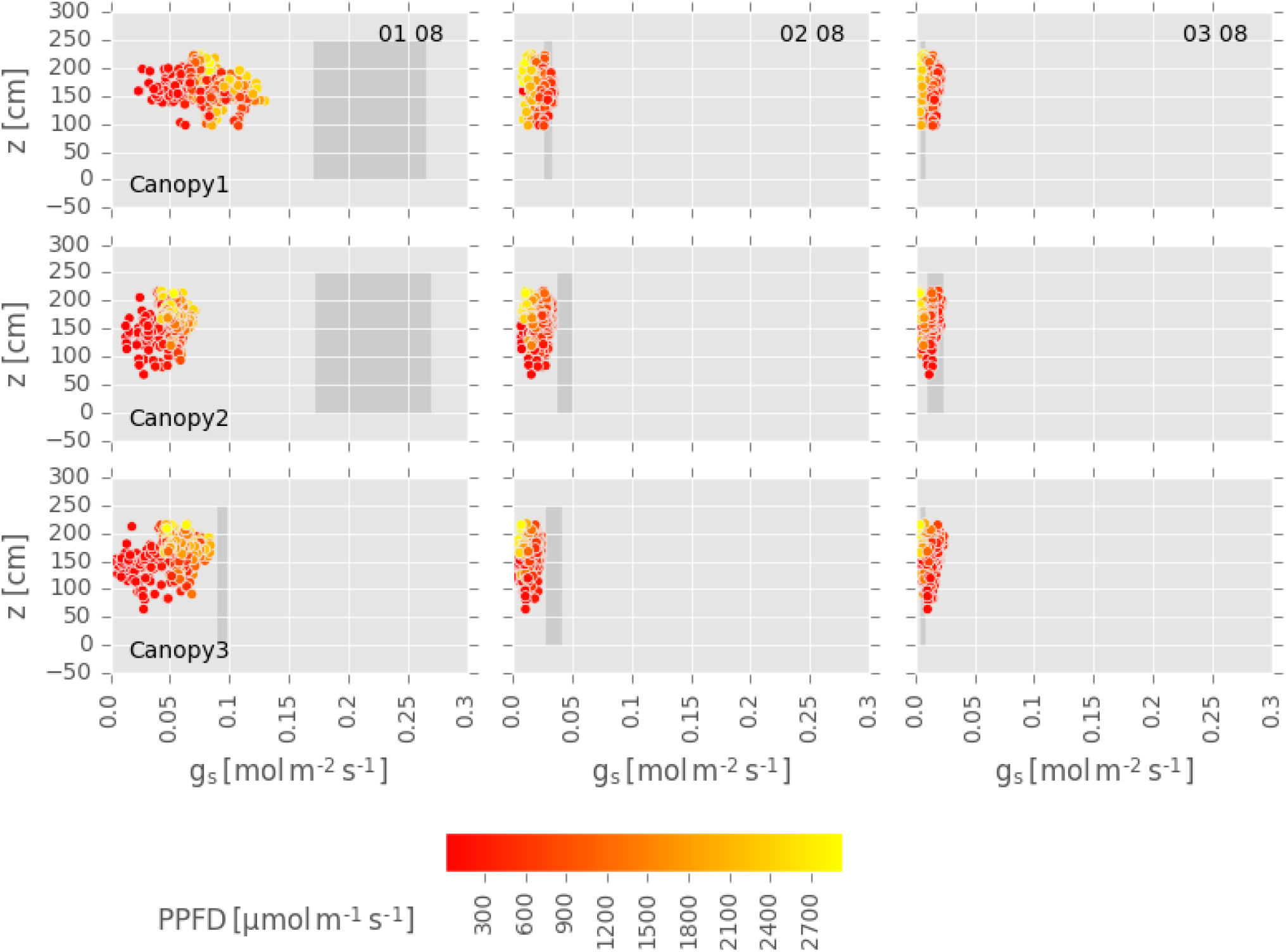
Snapshots of the stomatal conductance of the three GDC canopies considered in this study, referred to as ‘Canopy1’, ‘Canopy2’ and ‘Canopy3’, respectively, prior to solar noon, during the first three days following the onset of soil water deficit, respectively 01 08, 02 08 and 03 08. Soil predawn water potential of the three days was equal to −0.19, −0.38, −0.61 *MPa* respectively. Filled circles represent stomatal conductance to water (*g*_*s, H*_2_*O*_), their colors represent absorbed *PPFD* value per unlit leaf surface area; sunlit leaves are yellow while shaded leaves are red. Due to uncertainties in measurements locations, the observed stomatal conductance values of sunlit leaves are indicated by the dark grey patches which cover minimum and maximum values. Observed data were collected from experiments conducted in 2012.

Figures 9 and 10 show that the simulated *Ψ_leaf_* and *g*_*s, H*_2_*O*_ of the three GDC canopies decreased progressively as the soil water deficit increased, consistently with observations, but with an earlier onset of water stress which is probably due to inadequate parametrization either of the response function of *g*_*sH*_2_*O*_ to *Ψ_leaf_* (cf. Eq. 5b) or of the soil hydrodynamic model (cf. Coupling with irradiance and soil models). Upon the onset of water stress, when water deficit was still mild in the first day (date 01 08 in Figure 10), HydroShoot simulated higher *g*_*s, H*_2_*O*_ for sunlit leaves than for shaded leaves. Later, as water deficit increases, this trend is inverted, whereby sunlit leaves have the lowest *g*_*s, H*_2_*O*_ rates (dates 02 08 and 03 08 in Figure 10). This inversion is due to a lower *Ψ_leaf_* for sunlit leaves as a consequence of higher potential transpiration withdrawal per unit leaf surface area (cf. Eq. 1).

Moreover, Figure 10 shows that, by the end of the water-deficit period (day 03 08), leaf position had merely no more effect on its water vapor conductance *g*_*s, H*_2_*O*_. At this stage, water potential of all leaves reached low values at which stomata were almost closed. This uniformization of stomatal closure through the canopy is consistent with the observations reported by **Escalona et al. (2003, 2016)**. Both studies reported a progressive homogenization of gas-exchange rates of grapevine leaves (cv. Tempranillo, Manto Negro and Grenache) as soil water deficit increased. **Ngao et al. (2017)** reported similar results on apple trees (*Malus pumila* Mill.), showing that intra-canopy variability in *g*_*s, H*_2_*O*_ decreased significantly under the effect of soil water deficit.

The effect of soil water deficit on leaf temperature was efficiently captured by HydroShoot (Figure 11, Figure 12). The diurnal trends of leaves temperature were adequately reproduced (Figure 11) whereby a higher diurnal temperature was simulated under water-deficit (Figure 11b, d) than well-watered (Figure 11a, c) conditions. The comparison between simulated and observed temperatures at the leaf level (Figure 12) shows that the model simulated an increase in leaf-to-air temperature of approximately 2 *°C*, in agreement with observations (Figure 12a). Furthermore, HydroShoot reproduced the observed magnitude between minimum and maximum leaf temperatures across the canopy (Figure 12b) and how this magnitude increased with soil water deficit (Figure 12c), although underestimated the effect of water deficit on this magnitude (Figure 12c).

**Figure 11:**
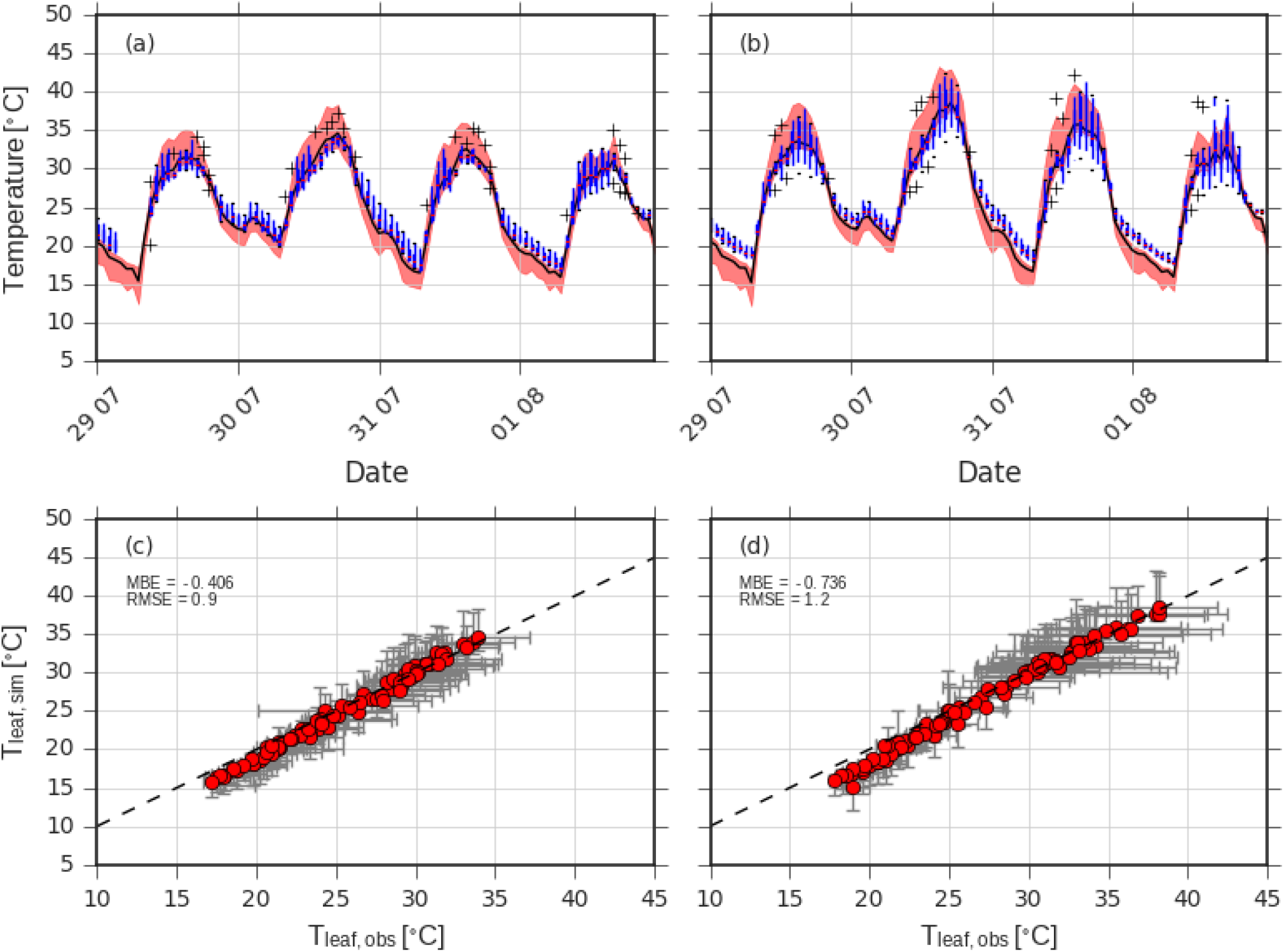
Comparison between simulated and observed individual leaf temperature for VSP canopies under well-watered (a and c) and water stressed (b and d) conditions. Soil predawn water potential of the four days was equal to −0.37, −0.50, −0.40, and −0.32 *MPa* respectively. (a and b) diurnal variation in leaf temperature: red zones indicate the extension between maximum and minimum simulated values, black curves indicate simulated mean values while blue boxplots indicate observed leaf temperature; (c and d) are 1:1 plots between observed (x-axis) and simulated (y-axis) leaves median temperatures for each hour with error bars representing minimum and maximum temperature values.

The overall adequation between simulated and observed leaf temperature was high when comparing median values (Figure 11) having an *RMSE* equal to 0.9 and 1.2 *°C* under well-watered and water-deficit conditions, respectively, while bias in *T_leaf_* was limited to a maximum value of −0.74 *°C*. However, greater error is obtained when leaf-to-leaf comparison is performed (Figure 12) with an overall *RMSE* reaching 2.3 *°C* and a bias of −0.79 *°C*. This is mainly due to the weak performance during nocturnal hours where temperature was underestimated by the model (Figure 11, Figure 12). This may originate either from the modelling approach or from the measurements. **Luquet et al. (2003)** reported similar trend of nocturnal temperature which they explained by the frequent dysfunctioning of thermocouples during the night. Similarly, **Bailey et al. (2016)** argued that such discrepancies originate rather from the thermocouples which may not measure precisely leaf temperature due to weak thermal conductivity between the leaf tissues and the wires. Thermocouples performance may explain in our study, at least partially, the discrepancies between observed and simulated leaf temperature during the night. We add, however, to this explanation, a second one which deals with the effect of the constant value used for sky emissivity that may have been inadequate knowing that the latter is affected by air humidity and temperature **(Brunt et al., 1932)**.

**Figure 12:**
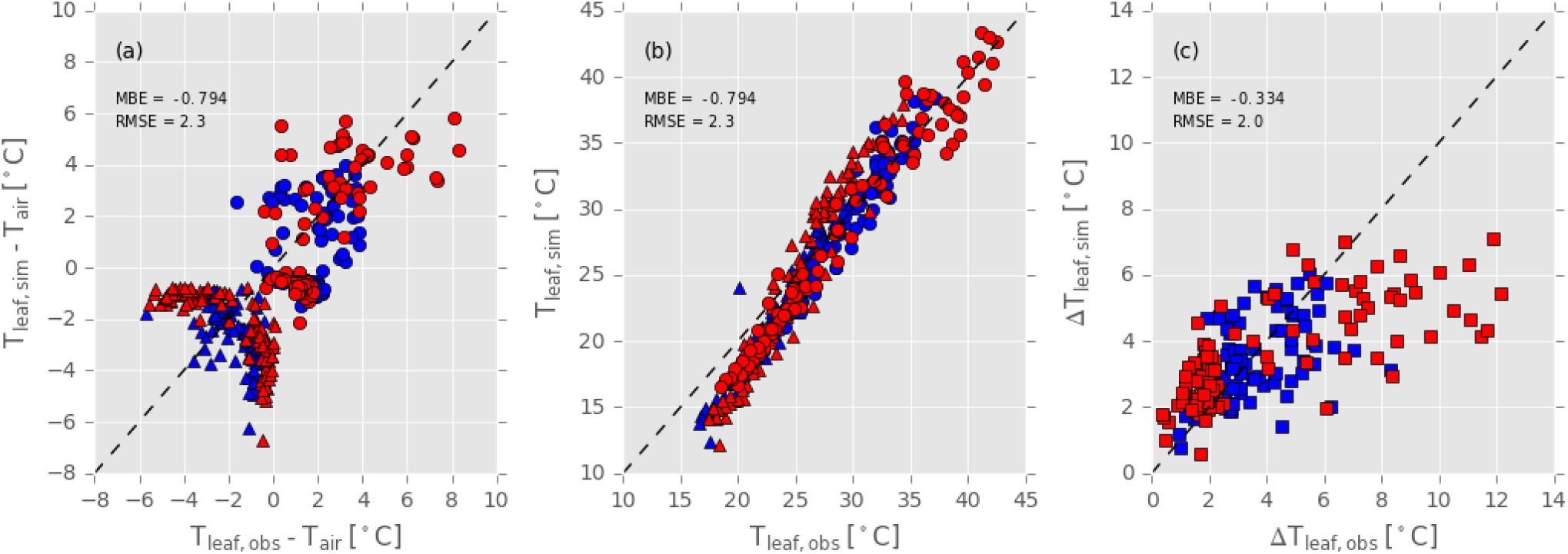
Comparison between the observed and simulated (a) differences between leaf and air temperatures, (b) leaf temperatures, and (c) magnitude between minimum and maximum leaf temperatures across the canopy of VSP plants. Maximum and minimum leaf temperatures during each hour are represented by circles and triangles, respectively (a and b plots), well-watered and water-deficit conditions are represented by blue red colors, respectively.

### Plant-scale

The observed daily patterns of *A_n, plant_* and *E_plant_* were accurately reproduced under both well-watered and water deficit conditions (Figure 13 for VSP and Figure 14 for GDC) albeit higher errors under water deficit occurred.

**Figure 13:**
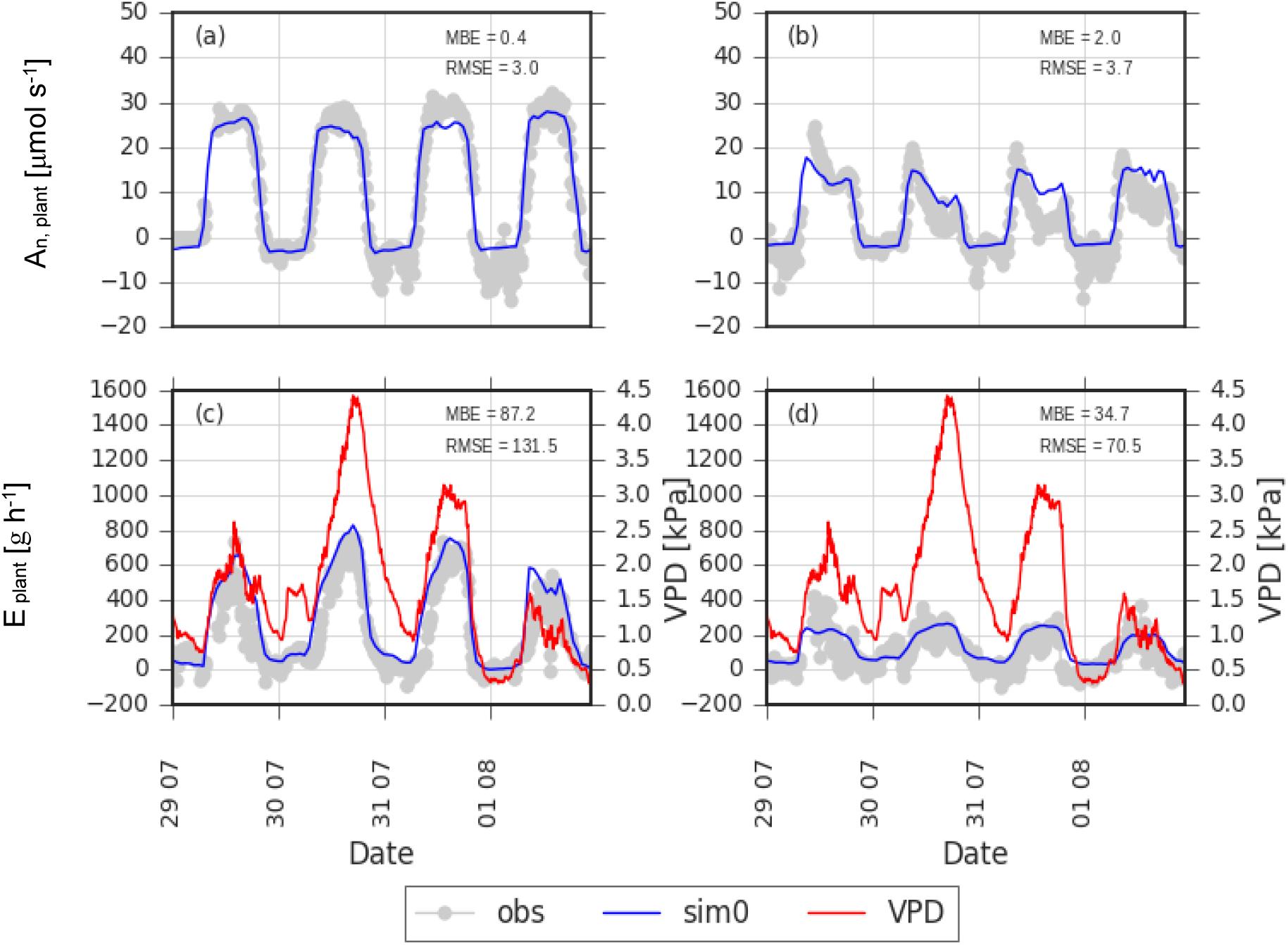
Comparison between the observed (grey circles) and simulated (blue circles) net carbon assimilation (*An, plant*) and transpiration (*E_plant_*) rates of entire grapevine plants (VSP training system) conducted under well-watered (left column) and water deficit (right column) conditions. Soil predawn water potential of the four days was equal to −0.37, −0.50, −0.40, and −0.32 *MPa* respectively. *MSE* and *RMSE* indicate respectively mean bias error and root mean squared error (same units of the y-axes). Red curves represent the vapor pressure deficit of the air. Observed data were collected from experiments conducted in 2009.

**Figure 14:**
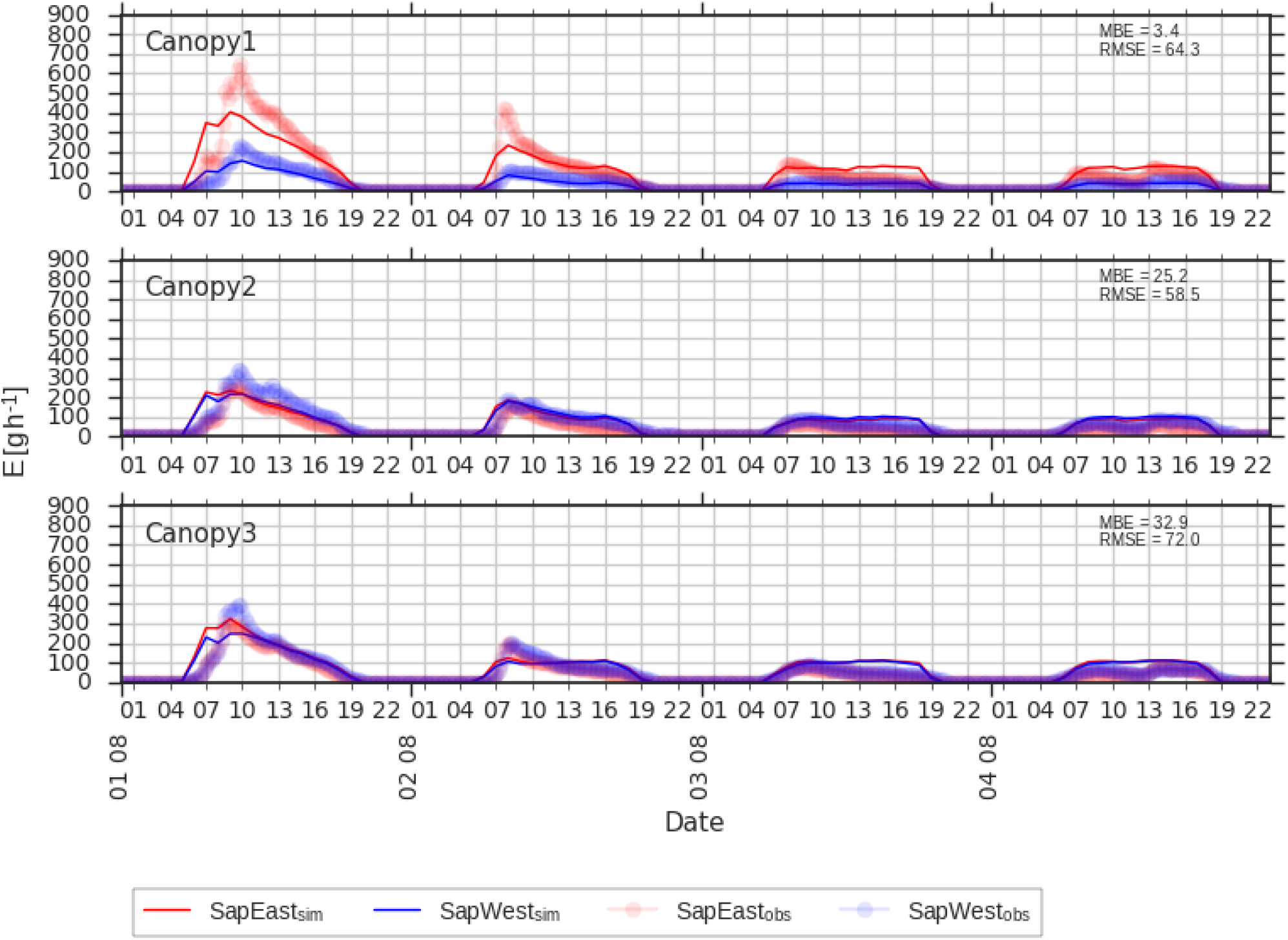
Comparison between the observed (circles) and simulated (curves) transpiration rates (*E_plant_*) for each cordon of three grapevines plants trained to GDC under water deficit conditions. Soil predawn water potential of the three days was equal to −0.19, −0.38, −0.61, and −0.51 *MPa* respectively. Red and blue colors indicate fluxes through east- and west-exposed cordons respectively. *MSE* and *RMSE* indicate respectively mean bias error and root mean squared error (same units of the y-axes).

For VSP canopies (Figure 13), the reduction in soil-water availability was reflected by the severe reductions in *A_n, plant_* and *E_plant_* rates, consistently with observations. *MBE* and *RMSE* totaled respectively 0.4 *μmol s*^−1^ and 3.0 *μmol s*^−1^ for *A_n, plant_* under well-watered conditions (Figure 13a), compared to 2.0 *μmol s*^−1^ and 3.7 *μmol s*^−1^, respectively, under water-deficit conditions (Figure 13b). Inversely, errors were lower for *E_plant_*, having *MBE* and *RMSE* at respectively 87.2 *g h*^−1^ and 131.5 *g h*^−1^ under well-watered conditions (Figure 13c), compared to 34.7 *g h*^−1^ and 70.5 *g h*^−1^, respectively under water-deficit conditions (Figure 13d).

For the three water-stressed GDC canopies (where only *E_plant_* rates were observed), HydroShoot captured reasonably the diurnal variations in *E_plant_* (Figure 14), having *MBE* falling between 3.4 and 32.9 *g h*^−1^ and *RMSE* between 58.5 and 72.0 *g h*^−1^ which were both similar to values obtained for VSP. It is noteworthy that the impact of the imbalance in leaf area between both cordons of Canopy1 (cf. Figure 4) was reflected in the simulated *E_plant_* (Figure 14, Canopy1) whereby a noticeable differences in the simulated fluxes was obtained between the eastern and western cordons (respectively red and blue curves in Figure 14, Canopy1) consistently with the observed sap flow rates (respectively red and blue dots in Figure 14, Canopy1). This example further demonstrate how the impact of canopy is adequately reflected on its ecophysiological functioning in HydroShoot.

### To which extent is modelling complexity needed?

We show in Figure 15 together with Table 1 that the best fit between simulated and observed gas-exchange rates was obtained using the complete HydroShoot model (i.e. sim0) which yielded, in almost all cases, the least values of *MBE* and *RMSE* for both *A_n, plant_* and *E_plant_*. However, the simulated hydraulic structure and spatialized leaf temperature values had unequal contributions to the predictions accuracy.

**Figure 15:**
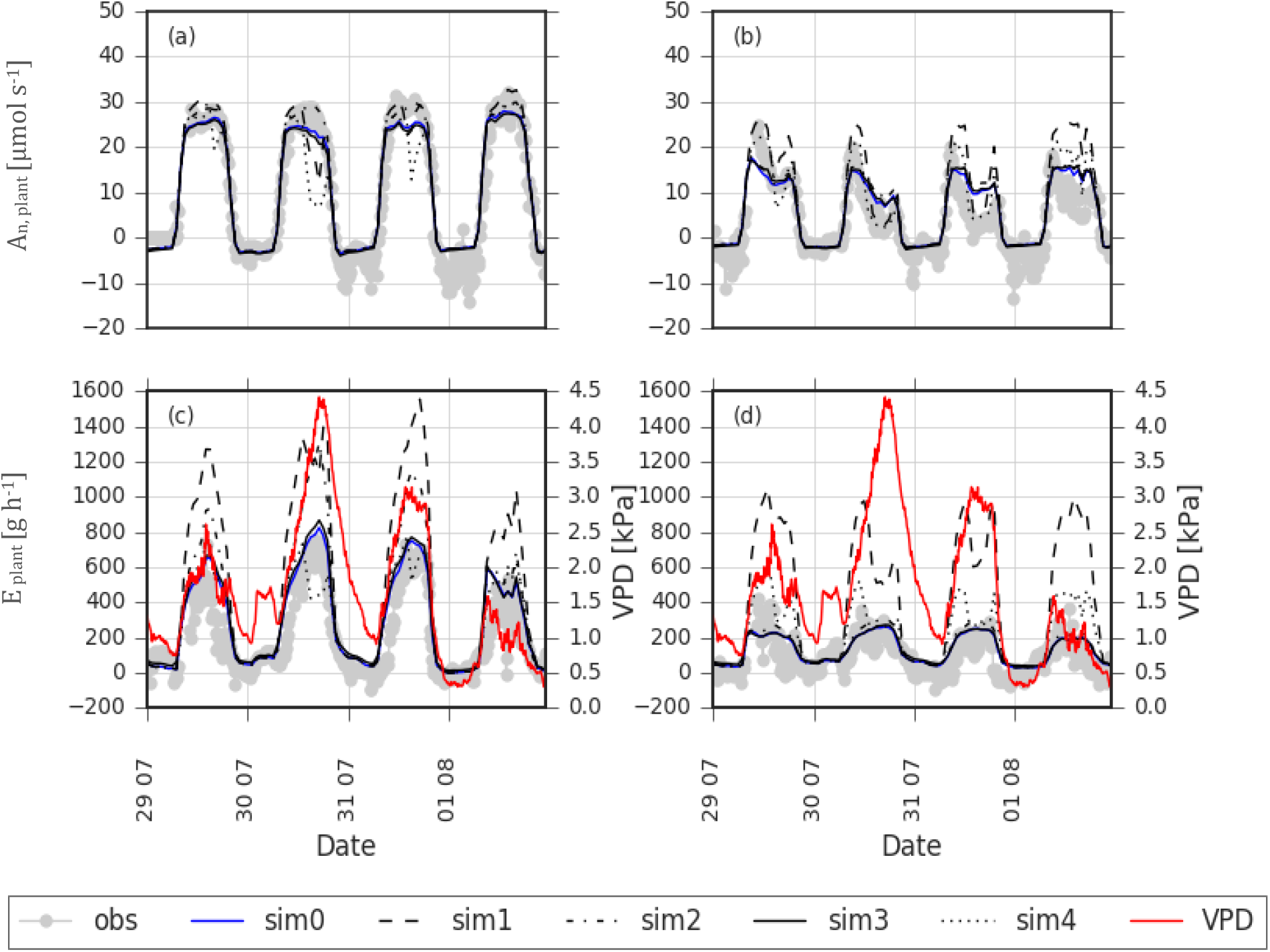
Impact of different modelling details of HydroShoot on the simulated plant carbon assimilation (*A_n, plant_*) and transpiration (*E_plant_*) rates: the reference (complete) HydroShoot version is indicated as sim0, sim1 indicates the version whereby leaf stomatal conductance varies with vapor pressure deficit instead of leaf water potential (i.e. original model of **Leuning, 1995**), sim2 indicates the results obtained when the shoot hydraulic structure was omitted (i.e. all leaves have the same water potential which is equal to that of the collar), sim3 is for results obtained by omitting energy balance of individual leaves (i.e. leaves temperature equal to air temperature) and sim4 is the same as sim1 but with an increased impact of the vapor pressure deficit on *g*_*s, CO*_2__ (*D*_0_ = 1 in Eq. 5a), finally, *obs* and *VPD* indicate respectively the observed gas rate and air vapor pressure deficit.

**Table 1:** Precision estimators of simulated carbon assimilation (*A_n, plant_*) and transpiration (*E_plant_*) rates of the plant of well-watered and water stressed VSP grapevines using five versions of HydroShoot: sim0, the reference (complete) HydroShoot version, sim1, the version whereby leaf stomatal conductance varies with vapor pressure deficit instead of leaf water potential (i.e. original model of Leuning, 1995), sim2, shoot hydraulic structure omitted (i.e. all leaves have the same water potential which is equal to that of the collar), sim3, energy balance of individual leaves omitted (i.e. leaves temperature equal to air temperature) and sim4, the same as sim1 but with an increased impact of the vapor pressure deficit on *g*_*s, CO*_2__ (*D*_0_ = 1 in Eq. 5a).

| | | $A_{n,plant}$ | | $E_{plant}$ | |
| --- | --- | --- | --- | --- | --- |
|  |  | MBE | RMSE | MBE | RMSE |
| | | $[\mu mol s^{-1}]$ | $[\mu mol s^{-1}]$ | $[g h^{-1}]$ | $[g h^{-1}]$ |
| WW | sim0 | 0.42 | 3.0 | 87.24 | 131.5 |
|  | sim1 | 1.46 | 4.0 | 304.37 | 433.1 |
|  | sim2 | 1.62 | 3.0 | 179.27 | 259.6 |
|  | sim3 | -0.02 | 3.1 | 108.04 | 149 |
|  | sim4 | -0.35 | 5.5 | 82.06 | 154.5 |
| WS | sim0 | 2.01 | 3.7 | 34.75 | 70.5 |
|  | sim1 | 4.58 | 6.9 | 319.08 | 441.0 |
|  | sim2 | 2.57 | 4.3 | 48.75 | 81.0 |
|  | sim3 | 2.13 | 3.9 | 40.99 | 74.0 |
|  | sim4 | 2.55 | 4.4 | 120.60 | 168.8 |

When stomatal aperture was dissociated from soil water status (i.e. sim1 and sim4), the results indicated a substantial increase in prediction errors, resulting in the highest values for both *MBE* and *RMSE* especially for *E_plant_*. For instance, *RMSE* increased from 131 to 433 *g h*^−1^ under well-watered conditions, and from 70 to 441 *g h*^−1^ under water-deficit conditions, when sim0 is compared to sim1. An improvement in prediction quality was obtained when stomatal aperture was linked to the collar water potential *Ψ_collar_* (i.e. sim2), yet, a considerable error still existed compared to the reference case. For instance, *MBE* of *A_n, plant_* increased from 0.42 to 1.62 *μmol s*^−1^ and from 2.01 to 2.51 *μmol s*^−1^, respectively under well-watered and water-deficit conditions, with sim2 compared to sim0. Similarly, *MBE* of *E_plant_* increased from 87 to 179 *g h*^−1^ and from 35 to 49 *g h*^−1^ respectively under well-watered and water-deficit conditions, using sim2 compared to the reference case sim0.

The aforementioned results indicate that linking leaf-scale stomatal aperture to leaf-level water potential (through *Ψ_leaf_*), by simulating the hydraulic structure, brought a considerable improvement to model performance. That is, predicting the intra-canopy variability in leaf water potential improves prediction accuracy of gas-exchange rates at the whole plant scale. These results agree with the conclusions reported by **Ngao et al. (2017)** on the role of leaf water potential variability on apple tree gas-exchange rates. The authors firstly reported that a reliable prediction of plant-scale gas-exchange fluxes in apple trees was allowed when stomatal closure was simulated as a function of soil water potential. However, they postulated that further improvements are yet expected when the hydraulic structure of the shoot is simulated.

Regarding the contribution of leaf-scale energy balance simulation on the predicted plant-scale fluxes, its effect was shown weak (Table 1). Indeed, disregarding energy balance calculations (i.e. sim3) did not affect, yet improved, the predicted *A_n, plant_* rates under well-watered conditions (*MBE* and *RMSE* changed from of 0.42 to −0.02 *μmol s*^−1^ and from 3.0 to 3.1 *μmol s*^−1^, respectively, using sim0 compared to sim3) and mildly affected *A_n, plant_* under water-deficit conditions (*MBE* and *RMSE* changed from of 2.01 to 2.13 *μmol s*^−1^ and from 3.7 to 3.9 *μmol s*^−1^, respectively, using sim0 compared to sim3). Similar results were also obtained for *E_plant_*.

Our results disagree with those reported by **Bauerle et al. (2007)** who estimated that disregarding the intra-canopy variability in leaf temperature would lead to overestimate *E_plant_* by 22 to 25% for Red Maple (*Acer rubrum* L.) In our case study on grapevine, disregarding leaf energy balance calculations increased *MBE* by 23% and 18% and *RMSE* by 13% and 5% for *E_plant_* under well-watered conditions and water-deficit conditions, respectively. The differences between our results and those reported by **Bauerle et al. (2007)** may rely on the way the authors accounted for the impact of microclimate inside the canopy on leaf photosynthetic traits. In their study, **Bauerle et al. (2007)** used leaf temperature as the primer driver for intra-canopy variability in leaf photosynthetic traits. In our study, we linked leaf photosynthetic traits to the 10-days cumulative absorbed *PPFD* (cf. Eq. 11) while leaf temperature was used only to affect directly *A_n_*, and indirectly *g*_*s, H*_2_*O*_. This conceptual difference may explain the lower sensitivity to leaf temperature in our study compared to that performed by **Bauerle et al. (2007)**. To this explanation may be added that provided by **Bailey et al. (2016)** who showed that the importance of accounting for the intra-canopy distribution of leaf temperature in FSPMs is a matter of canopy size. The authors reported that for grapevines, leaf temperature distribution had a negligible impact on the simulated plant-scale emitted thermal longwave irradiance. In contrast, on Freeman maple (*Acer* × *Freemanii*) which have notably higher leaf area per plant, simulating the spatial distribution of leaf temperature reduced by 50% prediction errors of the emitted thermal longwave irradiance. The low sensitivity of HydroShoot to leaf temperature may thus be linked to the simulated canopy size.

Notwithstanding, it is noteworthy that omitting energy balance calculations in HydroShoot allowed saving up to 21% calculation time on a portable computer. This considerable economy in calculation cost is an argument that should be considered when simulating large-scale plant scenes.

## Conclusions

We presented in this paper the functional-structural plant model (FSPM) HydroShoot. This model was built in order to allow simulating the effect of plant shoot architecture on its gas-exchange dynamics under soil water deficit conditions. In order to achieve this objective, we constructed HydroShoot on the base of three interacting processes: leaf-scale gas-exchange, leaf-scale energy balance, and internode-scale xylem transport (i.e. the hydraulic structure of the shoot). The produced model was evaluated using both virtual and real grapevine canopies of three strongly contrasting shoot architectures, under both well-watered and water-deficit conditions. We showed that HydroShoot reproduced efficiently the effect of canopy architecture on plant-scale gas-exchange processes under the observed gradient of water deficit conditions, fulfilling thus the objectives for which the model was built. We showed furthermore that both hydraulic structure and energy balance simulations were required for a precise prediction of plant-scale gas-exchange rates under soil water deficit. However, our results indicate that the hydraulic structure has, by far, the largest effect on simulated net photosynthesis and transpiration rates for grapevine. In contrast, simulating leaf-scale energy balance improves minorly prediction results.

## Acknowledgments

This research was funded by the European Community’s Seventh Framework Program (FP7/2007-2013) under the grant agreement no. FP7-311775, Project INNOVINE. It was also partly funded by the Environment and Agronomy department of the French National Institute for Agricultural Research (INRA).

The authors greatly acknowledge Mr. Gerardo Lopez for his concise and constructive notes on this manuscript.

The authors dedicate this work to their beloved colleague Eric Lebon, who left us just before the closure of the INNOVINE project.

## Appendices

### I. Equations of the net CO_2_ assimilation sub-model

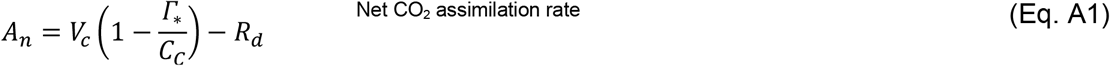

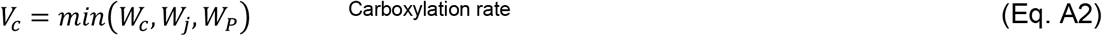

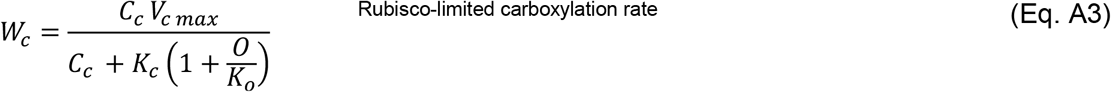

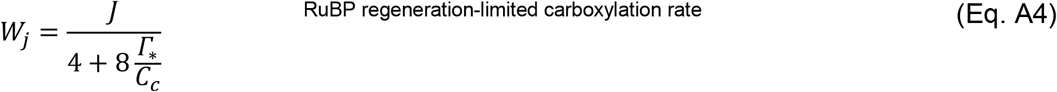

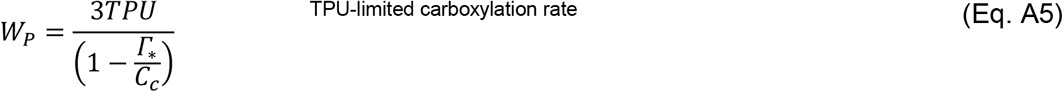

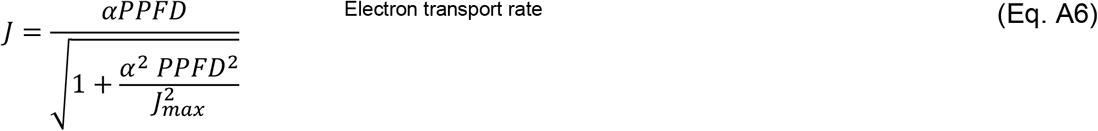

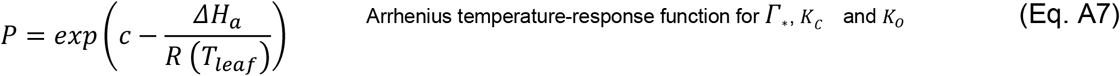

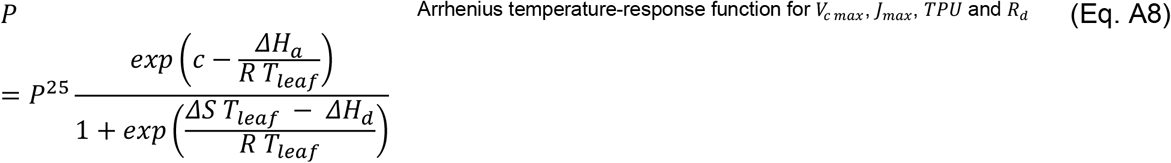

### II. Empirical photoinhibition model

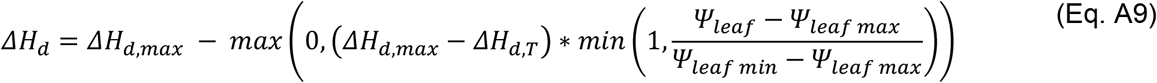

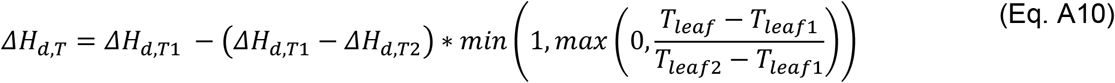

where *ΔH_d_* [*kJ mol*_*CO*_2__^−1^] is calculated after accounting for the joint effects of leaf water potential *Ψ_leaf_* [*MPa*] and temperature *T_leaf_* [*K*], *ΔH_d, max_* [*kJ mol*_*CO*_2__ ^−1^] is the value of *ΔH_d_* without accounting for photoinhibition; *ΔH_d, T_* [*kJ mol*_*CO*_2__^−1^] is the value of *ΔH_d_* after accounting for the effect of *T_leaf_*; *Ψ_leaf max_* and *Ψ_leaf min_* [*MPa*] are leaf water potential values at which photoinhibition starts and reaches its maximum effect, respectively; finally, *ΔH*_*d, T*1_ and *ΔH*_*d, T*2_ [*kJ mol*_*CO*_2__^−1^] are empirical thresholds corresponding to leaf temperatures *T*_*leaf*1_ and *T*_*leaf*2_ which are temperatures at which photoinhibition starts and reaches its maximum effect, respectively.

### III. Numerical resolution

#### Initialization

i. Photosynthetic capacity parameters of individual leaves are calculated based on *PPFD_10_* values using equations 9 to 11;
ii. Leaf temperature (*T_i_*) is assumed equal to that of the previous time step or equal to air temperature (*T_air_*) for the first time step.
iii. Collar water potential *Ψ_collar_* is forced equal to soil water potential *Ψ_soil_* as lower boundary of the hydraulic structure (only if the option of hydraulic structure is considered);
iv. Xylem maximum and actual hydraulic conductivities of each internode (respectively *K_max, i_* and *K_i_*) are calculated using respectively equations 2 and 3;
v. Xylem water potential at each node (*Ψ_u, i_*) is assumed equal to that of the previous time-step, otherwise, it is calculated using Eq. 1, assuming maximum hydraulic conductivities in all plant segments and considering the plant to be in hydrostatic equilibrium with the soil.

#### Convergence

When Eq. 1 is applied to a plant shoot of *N* segments, thus consisting of *N* nodes according to the graph theory, a system of *N* equations for *N + 1* water head values is obtained. To solve the system for water potentials at all nodes, boundary conditions are set as follows: the lower boundary condition is a Dirichlet-type whereby a constant soil water potential value (*Ψ_soil_*) is forced during each iteration; the upper boundary condition is a Neumann-type, whereby a constant flow (*F_i_*) is forced at each leaf node (*F_i_* is equal to transpiration *E* multiplied by leaf surface, cf. Eq. 6). The calculation procedure is detailed in the numerical resolution section.
1. Leaf-scale *A_n_* and *E* rates are calculated using equations A1 to A8 and 4 to 8;
2. *E* is multiplied by leaf surface area to obtain xylem flux (*F_i_*) at the leaf level, that is forced to the hydraulic structure system;
3. *F_i_* values are calculated for each internode by “walking downwards” from leaves to collar (postorder traversal on MTG);
4. New values of *Ψ_u, i_* are calculated from Eq. 1, assuming constant *K_i_*, by “walking upwards” (preorder traversal on MTG) starting from the collar (where *Ψ_collar_* is known) up to the leaves;
5. *K_i_* values are updated using Eq. 2;
6. Steps 4 and 5 are repeated until convergence, that is the absolute difference (*ΔΨ_u, i_*) between two consecutive iterations (respectively *j – 1* and *j*) of at most one node, is lower than a predefined error threshold (*ε_x_*):

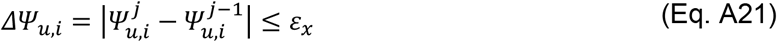
7. New *T_i_* values are calculated from energy balance, considering the new *E* values in Eq. 13 using the following steps;
  a. In a first step, the surrounding temperature (*T_leaves_*) is fixed at the first iteration (equal to *T_leaves_* from the previous calculation step) and *T_i_* is solved for each leaf.
  b. In a second step, *T_leaves_* is updated considering the new values of leaf temperature and a new set of *T_i_* values is computed.
  c. The first and second steps are repeated until the maximum absolute difference of the temperature of each leaf between two consecutive iterations falls below 0.02 °C **(Maes and Steppe, 2012)**.
8. Steps 1 to 7 are repeated until convergence, that is the absolute difference (*ΔT_i_*) between two consecutive iterations (respectively *j – 1* and *j*) of at most one leaf, is lower than a predefined error threshold (*ε_T_*):

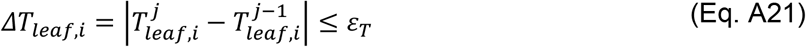

### IV. Table of variable symbols, values and units used for CO_2_ net assimilation, stomatal conductance and hydraulic structure submodels

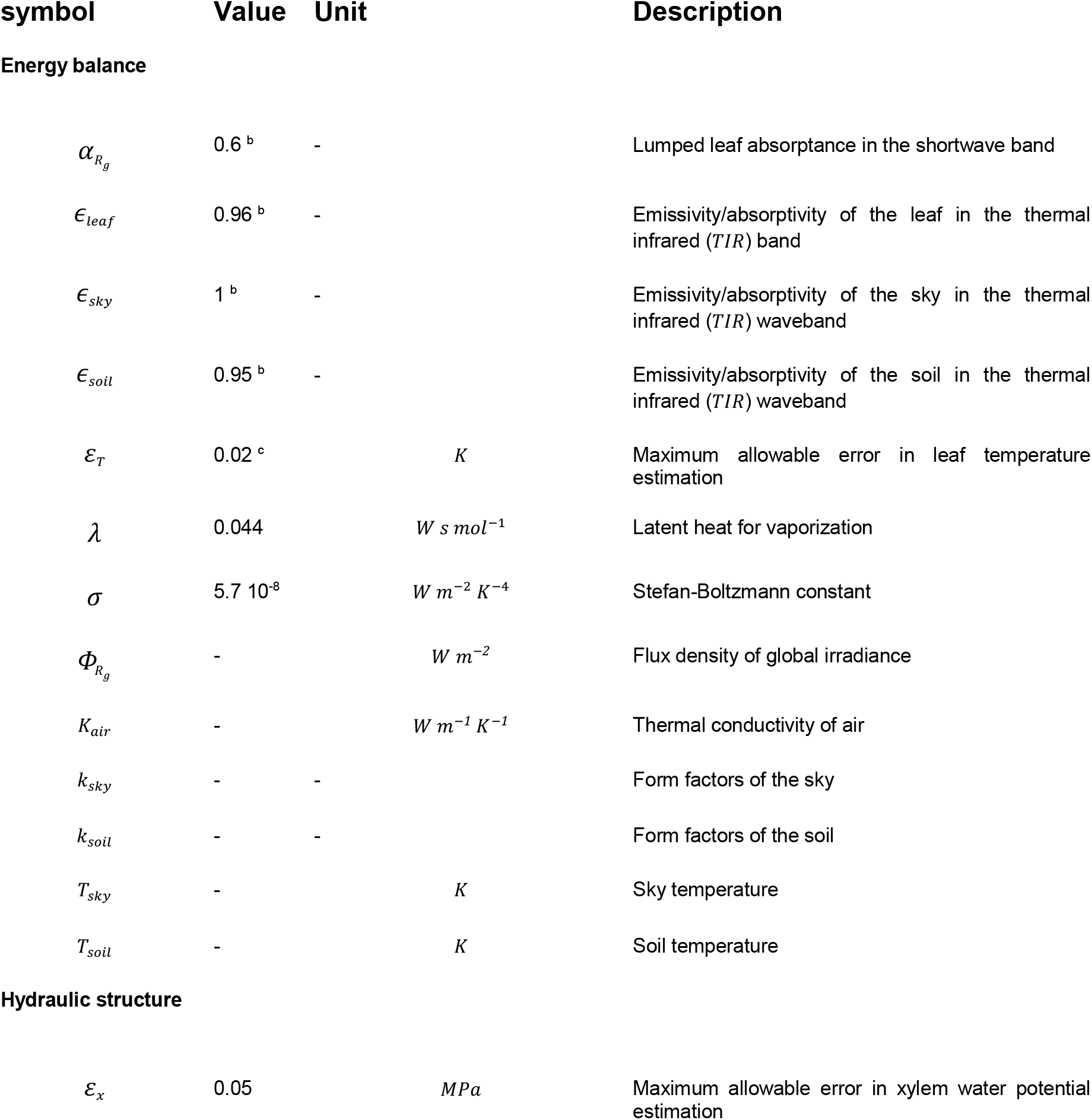

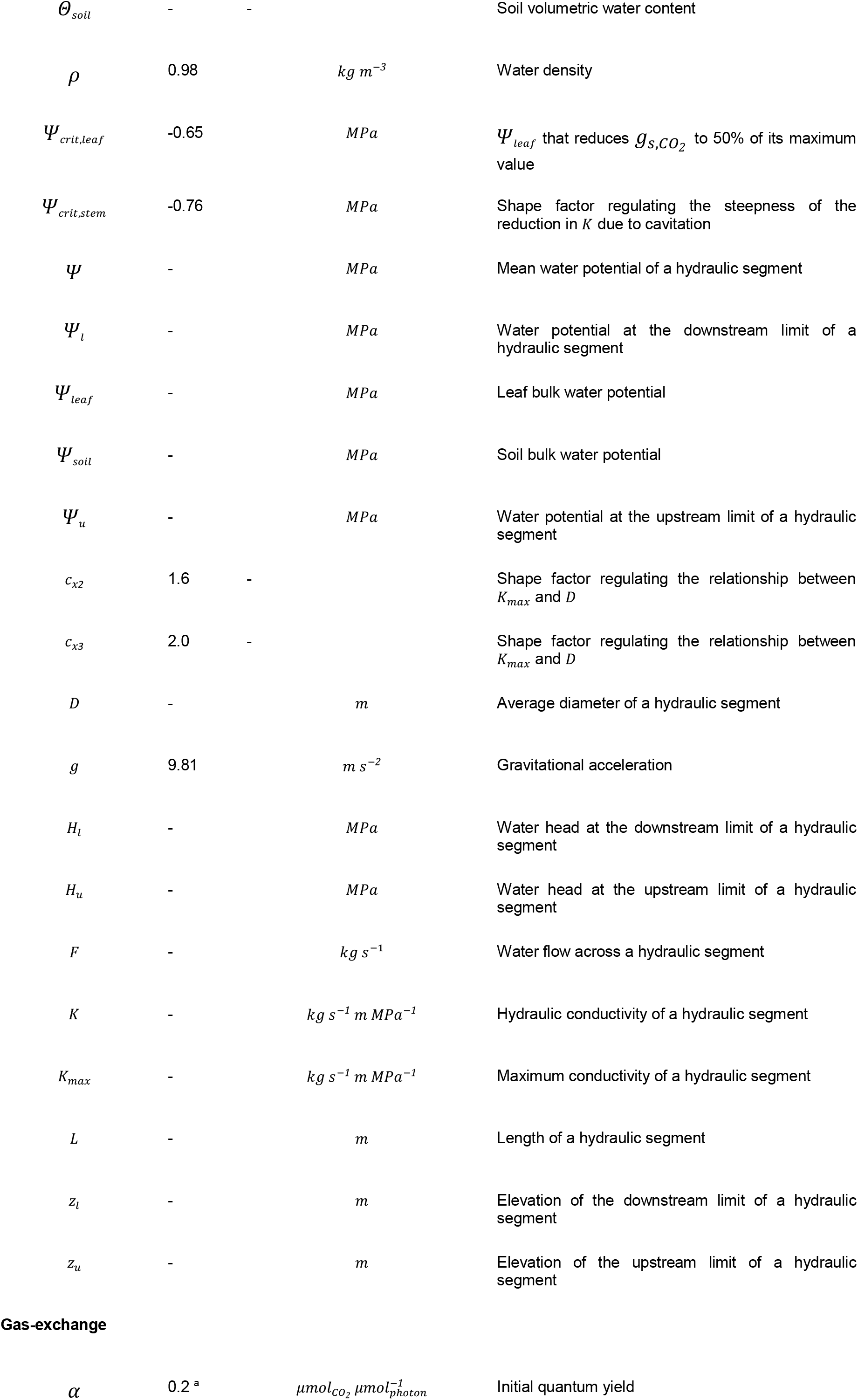

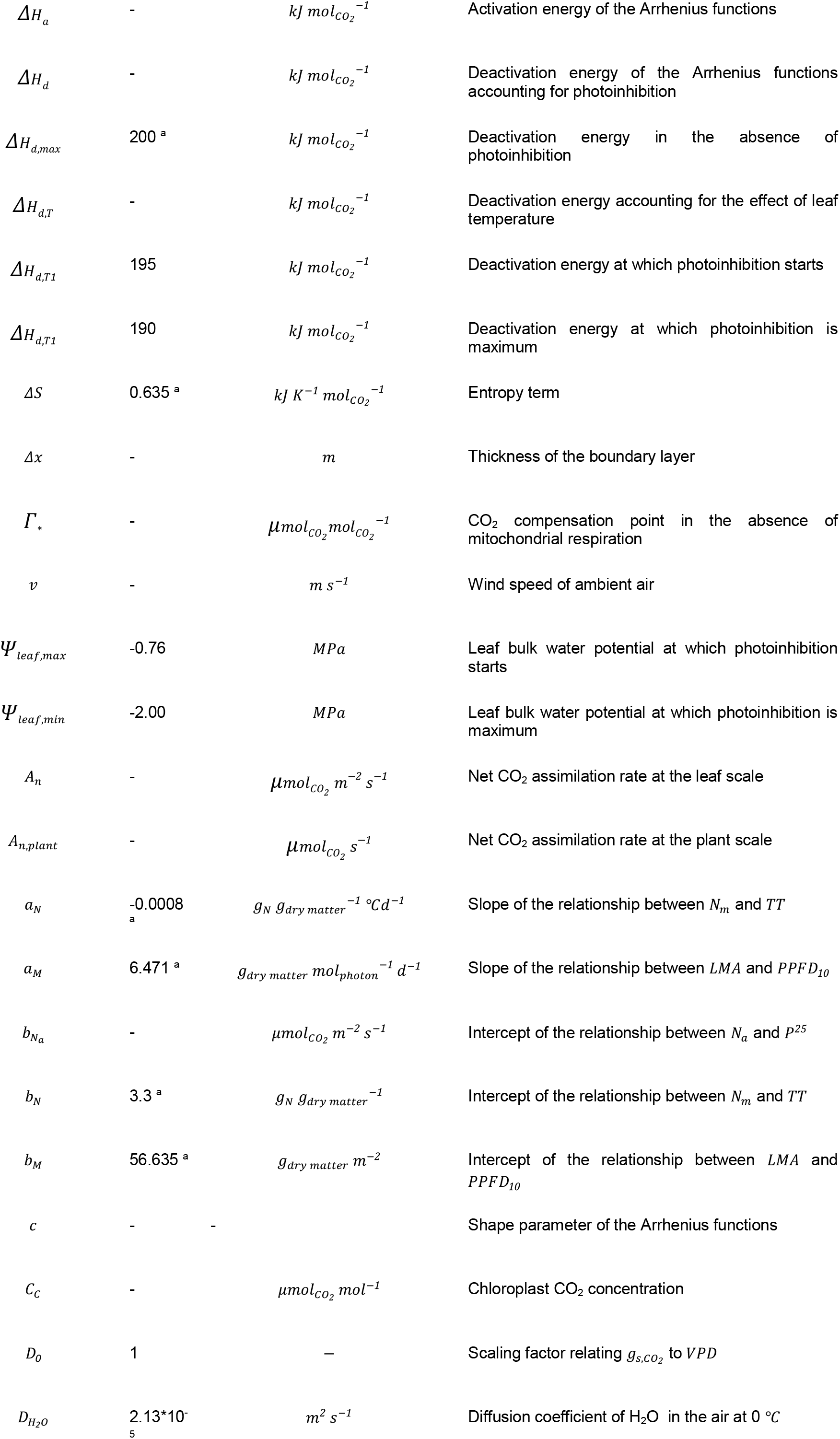

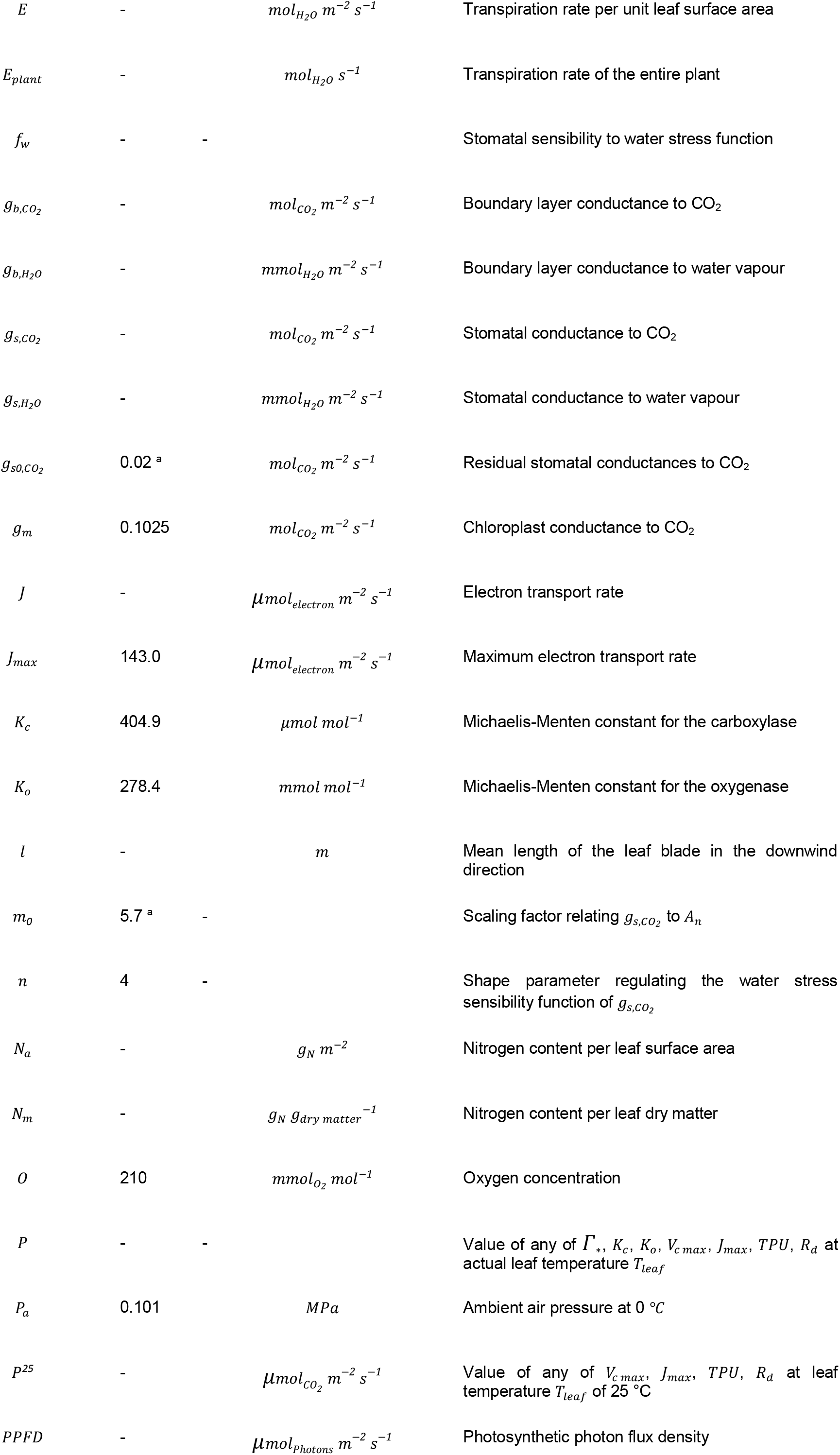

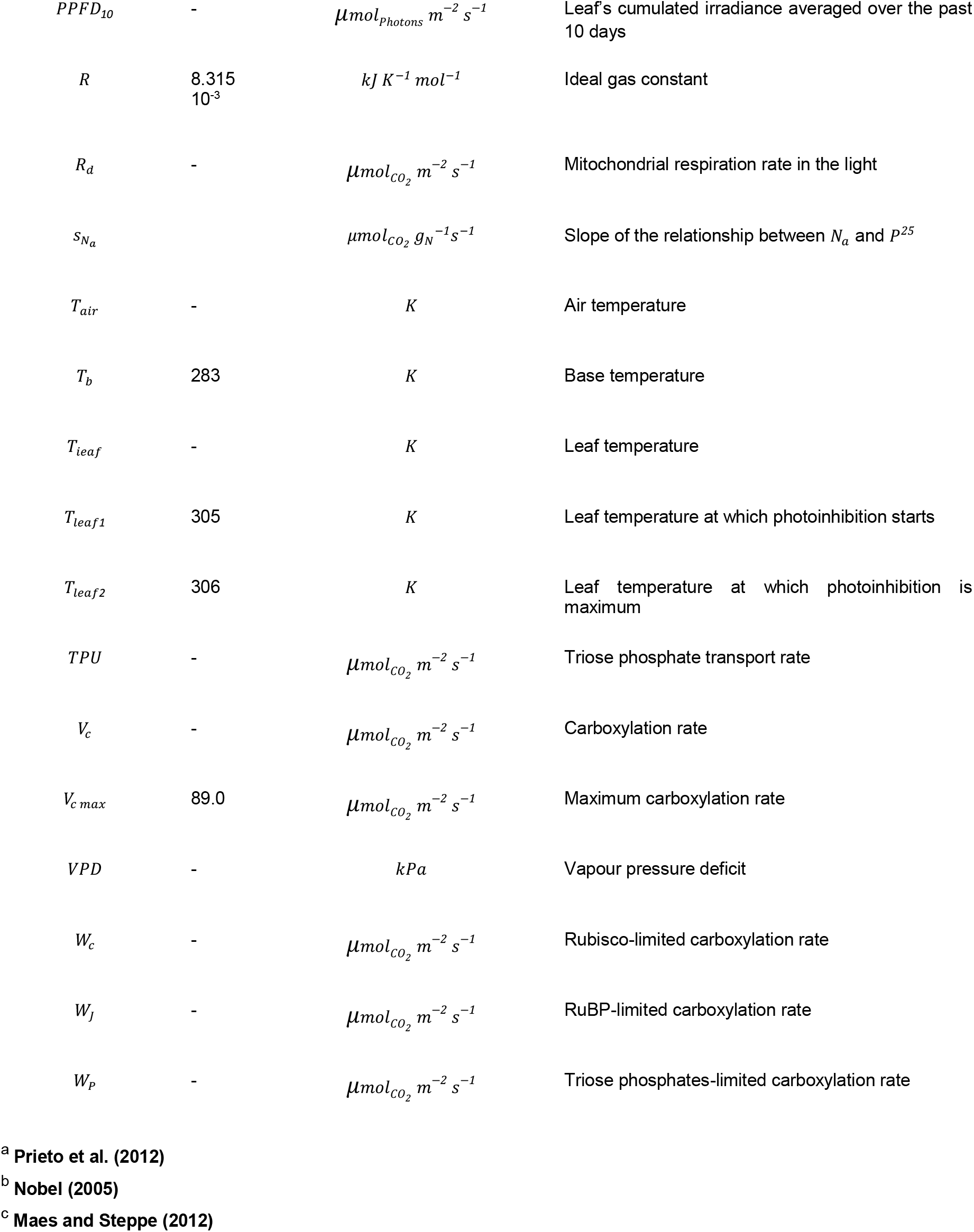

IV. Table of variable symbols, values and units used for CO<sub>2</sub> net assimilation, stomatal conductance and hydraulic structure submodels.
| symbol | Value | Unit | Description |
| --- | --- | --- | --- |
| **Energy balance** |  |  |  |
| $\alpha_{R_g}$ | 0.6<sup>b</sup> | - | Lumped leaf absorptance in the shortwave band |
| $\epsilon_{leaf}$ | 0.96<sup>b</sup> | - | Emissivity/absorptivity of the leaf in the thermal infrared (*TIR*) band |
| $\epsilon_{sky}$ | 1<sup>b</sup> | - | Emissivity/absorptivity of the sky in the thermal infrared (*TIR*) waveband |
| $\epsilon_{soil}$ | 0.95<sup>b</sup> | - | Emissivity/absorptivity of the soil in the thermal infrared (*TIR*) waveband |
| $\varepsilon_T$ | 0.02<sup>c</sup> | $K$ | Maximum allowable error in leaf temperature estimation |
| $\lambda$ | 0.044 | $W\ s\ mol^{-1}$ | Latent heat for vaporization |
| $\sigma$ | $5.7\ 10^{-8}$ | $W\ m^{-2}\ K^{-4}$ | Stefan-Boltzmann constant |
| $\Phi_{R_g}$ | - | $W\ m^{-2}$ | Flux density of global irradiance |
| $K_{air}$ | - | $W\ m^{-1}\ K^{-1}$ | Thermal conductivity of air |
| $k_{sky}$ | - | - | Form factors of the sky |
| $k_{soil}$ | - | - | Form factors of the soil |
| $T_{sky}$ | - | $K$ | Sky temperature |
| $T_{soil}$ | - | $K$ | Soil temperature |
| **Hydraulic structure** |  |  |  |
| $\varepsilon_x$ | 0.05 | $MPa$ | Maximum allowable error in xylem water potential estimation |

|  |  |  |  |  |
| --- | --- | --- | --- | --- |
| $\theta_{soil}$ | - | - | | Soil volumetric water content |
| $\rho$ | 0.98 | | $kg\ m^{-3}$ | Water density |
| $\Psi_{crit,leaf}$ | -0.65 | | $MPa$ | $\Psi_{leaf}$ that reduces $g_{S,CO_2}$ to 50% of its maximum value |
| $\Psi_{crit,stem}$ | -0.76 | | $MPa$ | Shape factor regulating the steepness of the reduction in $K$ due to cavitation |
| $\psi$ | - | | $MPa$ | Mean water potential of a hydraulic segment |
| $\psi_l$ | - | | $MPa$ | Water potential at the downstream limit of a hydraulic segment |
| $\Psi_{leaf}$ | - | | $MPa$ | Leaf bulk water potential |
| $\Psi_{soil}$ | - | | $MPa$ | Soil bulk water potential |
| $\Psi_u$ | - | | $MPa$ | Water potential at the upstream limit of a hydraulic segment |
| $c_{x2}$ | 1.6 | - | | Shape factor regulating the relationship between $K_{max}$ and $D$ |
| $c_{x3}$ | 2.0 | - | | Shape factor regulating the relationship between $K_{max}$ and $D$ |
| $D$ | - | | $m$ | Average diameter of a hydraulic segment |
| $g$ | 9.81 | | $m\ s^{-2}$ | Gravitational acceleration |
| $H_l$ | - | | $MPa$ | Water head at the downstream limit of a hydraulic segment |
| $H_u$ | - | | $MPa$ | Water head at the upstream limit of a hydraulic segment |
| $F$ | - | | $kg\ s^{-1}$ | Water flow across a hydraulic segment |
| $K$ | - | | $kg\ s^{-1}\ m\ MPa^{-1}$ | Hydraulic conductivity of a hydraulic segment |
| $K_{max}$ | - | | $kg\ s^{-1}\ m\ MPa^{-1}$ | Maximum conductivity of a hydraulic segment |
| $L$ | - | | $m$ | Length of a hydraulic segment |
| $z_l$ | - | | $m$ | Elevation of the downstream limit of a hydraulic segment |
| $z_u$ | - | | $m$ | Elevation of the upstream limit of a hydraulic segment |

Gas-exchange
|  |  |  |  |  |
| --- | --- | --- | --- | --- |
| $\alpha$ | 0.2<sup>a</sup> | | $\mu mol_{CO_2}\ \mu mol_{photon}^{-1}$ | Initial quantum yield |
| $\Delta H_a$ | - | $\text{kJ mol}_{\text{CO}_2}^{-1}$ | Activation energy of the Arrhenius functions |
| $\Delta H_d$ | - | $\text{kJ mol}_{\text{CO}_2}^{-1}$ | Deactivation energy of the Arrhenius functions accounting for photoinhibition |
| $\Delta H_{d,max}$ | 200<sup>a</sup> | $\text{kJ mol}_{\text{CO}_2}^{-1}$ | Deactivation energy in the absence of photoinhibition |
| $\Delta H_{d,T}$ | - | $\text{kJ mol}_{\text{CO}_2}^{-1}$ | Deactivation energy accounting for the effect of leaf temperature |
| $\Delta H_{d,T1}$ | 195 | $\text{kJ mol}_{\text{CO}_2}^{-1}$ | Deactivation energy at which photoinhibition starts |
| $\Delta H_{d,T1}$ | 190 | $\text{kJ mol}_{\text{CO}_2}^{-1}$ | Deactivation energy at which photoinhibition is maximum |
| $\Delta S$ | 0.635<sup>a</sup> | $\text{kJ K}^{-1} \text{mol}_{\text{CO}_2}^{-1}$ | Entropy term |
| $\Delta x$ | - | $m$ | Thickness of the boundary layer |
| $\Gamma_*$ | - | $\mu\text{mol}_{\text{CO}_2} \text{mol}_{\text{CO}_2}^{-1}$ | CO<sub>2</sub> compensation point in the absence of mitochondrial respiration |
| $v$ | - | $\text{m s}^{-1}$ | Wind speed of ambient air |
| $\Psi_{leaf,max}$ | -0.76 | $\text{MPa}$ | Leaf bulk water potential at which photoinhibition starts |
| $\Psi_{leaf,min}$ | -2.00 | $\text{MPa}$ | Leaf bulk water potential at which photoinhibition is maximum |
| $A_n$ | - | $\mu\text{mol}_{\text{CO}_2} \text{m}^{-2} \text{s}^{-1}$ | Net CO<sub>2</sub> assimilation rate at the leaf scale |
| $A_{n,plant}$ | - | $\mu\text{mol}_{\text{CO}_2} \text{s}^{-1}$ | Net CO<sub>2</sub> assimilation rate at the plant scale |
| $a_N$ | -0.0008<sup>a</sup> | $g_N g_{dry\ matter}^{-1} ^\circ\text{C d}^{-1}$ | Slope of the relationship between $N_m$ and $TT$ |
| $a_M$ | 6.471<sup>a</sup> | $g_{dry\ matter} \text{mol}_{\text{photon}}^{-1} \text{d}^{-1}$ | Slope of the relationship between $LMA$ and $PPFD_{10}$ |
| $b_{N_a}$ | - | $\mu\text{mol}_{\text{CO}_2} \text{m}^{-2} \text{s}^{-1}$ | Intercept of the relationship between $N_a$ and $P^{25}$ |
| $b_N$ | 3.3<sup>a</sup> | $g_N g_{dry\ matter}^{-1}$ | Intercept of the relationship between $N_m$ and $TT$ |
| $b_M$ | 56.635<sup>a</sup> | $g_{dry\ matter} \text{m}^{-2}$ | Intercept of the relationship between $LMA$ and $PPFD_{10}$ |
| $c$ | - | - | Shape parameter of the Arrhenius functions |
| $C_C$ | - | $\mu\text{mol}_{\text{CO}_2} \text{mol}^{-1}$ | Chloroplast CO<sub>2</sub> concentration |
| $D_0$ | 1 | — | Scaling factor relating $g_{s,\text{CO}_2}$ to $VPD$ |
| $D_{H_2O}$ | 2.13*10<sup>-5</sup> | $\text{m}^2 \text{s}^{-1}$ | Diffusion coefficient of H<sub>2</sub>O in the air at 0 °C |

|  |  |  |  |  |
| --- | --- | --- | --- | --- |
| $E$ | - | | $mol_{H_2O} m^{-2} s^{-1}$ | Transpiration rate per unit leaf surface area |
| $E_{plant}$ | - | | $mol_{H_2O} s^{-1}$ | Transpiration rate of the entire plant |
| $f_w$ | - | - | | Stomatal sensibility to water stress function |
| $g_{b,CO_2}$ | - | | $mol_{CO_2} m^{-2} s^{-1}$ | Boundary layer conductance to CO<sub>2</sub> |
| $g_{b,H_2O}$ | - | | $mmol_{H_2O} m^{-2} s^{-1}$ | Boundary layer conductance to water vapour |
| $g_{s,CO_2}$ | - | | $mol_{CO_2} m^{-2} s^{-1}$ | Stomatal conductance to CO<sub>2</sub> |
| $g_{s,H_2O}$ | - | | $mmol_{H_2O} m^{-2} s^{-1}$ | Stomatal conductance to water vapour |
| $g_{s0,CO_2}$ | 0.02<sup>a</sup> | | $mol_{CO_2} m^{-2} s^{-1}$ | Residual stomatal conductances to CO<sub>2</sub> |
| $g_m$ | 0.1025 | | $mol_{CO_2} m^{-2} s^{-1}$ | Chloroplast conductance to CO<sub>2</sub> |
| $J$ | - | | $\mu mol_{electron} m^{-2} s^{-1}$ | Electron transport rate |
| $J_{max}$ | 143.0 | | $\mu mol_{electron} m^{-2} s^{-1}$ | Maximum electron transport rate |
| $K_c$ | 404.9 | | $\mu mol mol^{-1}$ | Michaelis-Menten constant for the carboxylase |
| $K_o$ | 278.4 | | $mmol mol^{-1}$ | Michaelis-Menten constant for the oxygenase |
| $l$ | - | | $m$ | Mean length of the leaf blade in the downwind direction |
| $m_0$ | 5.7<sup>a</sup> | - | | Scaling factor relating $g_{s,CO_2}$ to $A_n$ |
| $n$ | 4 | - | | Shape parameter regulating the water stress sensibility function of $g_{s,CO_2}$ |
| $N_a$ | - | | $g_N m^{-2}$ | Nitrogen content per leaf surface area |
| $N_m$ | - | | $g_N g_{dry matter}^{-1}$ | Nitrogen content per leaf dry matter |
| $O$ | 210 | | $mmol_{O_2} mol^{-1}$ | Oxygen concentration |
| $P$ | - | - | | Value of any of $\Gamma^*$, $K_c$, $K_o$, $V_{c max}$, $J_{max}$, $TPU$, $R_d$ at actual leaf temperature $T_{leaf}$ |
| $P_a$ | 0.101 | | $MPa$ | Ambient air pressure at 0 °C |
| $p^{25}$ | - | | $\mu mol_{CO_2} m^{-2} s^{-1}$ | Value of any of $V_{c max}$, $J_{max}$, $TPU$, $R_d$ at leaf temperature $T_{leaf}$ of 25 °C |
| $PPFD$ | - | | $\mu mol_{photons} m^{-2} s^{-1}$ | Photosynthetic photon flux density |
| $PPFD_{10}$ | - | $\mu mol_{photons} m^{-2} s^{-1}$ | Leaf's cumulated irradiance averaged over the past 10 days |
| $R$ | 8.315<br>$10^{-3}$ | $kJ K^{-1} mol^{-1}$ | Ideal gas constant |
| $R_d$ | - | $\mu mol_{CO_2} m^{-2} s^{-1}$ | Mitochondrial respiration rate in the light |
| $s_{N_a}$ | | $\mu mol_{CO_2} g_N^{-1} s^{-1}$ | Slope of the relationship between $N_a$ and $P^{25}$ |
| $T_{air}$ | - | $K$ | Air temperature |
| $T_b$ | 283 | $K$ | Base temperature |
| $T_{leaf}$ | - | $K$ | Leaf temperature |
| $T_{leaf1}$ | 305 | $K$ | Leaf temperature at which photoinhibition starts |
| $T_{leaf2}$ | 306 | $K$ | Leaf temperature at which photoinhibition is maximum |
| $TPU$ | - | $\mu mol_{CO_2} m^{-2} s^{-1}$ | Triose phosphate transport rate |
| $V_c$ | - | $\mu mol_{CO_2} m^{-2} s^{-1}$ | Carboxylation rate |
| $V_{c\ max}$ | 89.0 | $\mu mol_{CO_2} m^{-2} s^{-1}$ | Maximum carboxylation rate |
| $VPD$ | - | $kPa$ | Vapour pressure deficit |
| $W_c$ | - | $\mu mol_{CO_2} m^{-2} s^{-1}$ | Rubisco-limited carboxylation rate |
| $W_J$ | - | $\mu mol_{CO_2} m^{-2} s^{-1}$ | RuBP-limited carboxylation rate |
| $W_P$ | - | $\mu mol_{CO_2} m^{-2} s^{-1}$ | Triose phosphates-limited carboxylation rate |

<sup>a</sup> Prieto et al. (2012)
<sup>b</sup> Nobel (2005)
<sup>c</sup> Maes and Steppe (2012)

## Supplementary material

### Data collection from experiments

Data for model evaluation were collected from experiments conducted in 2009 and 2012 on grapevine (cv. Syrah, grafted on SO4) at INRA, in Montpellier (3°53” E, 43°37” N, 44 m alt). 5 grapevines trained with two contrasting training systems were considered (cf. Figure 5 in the paper): GDC in 2009 and VSP in 2012. Grapevine rows were oriented 140° from North on a shallow sandy loam soil with a low water holding capacity. Inter-row spacing was 3.6 *m* for GDC and 1.8 *m* for VSP. Intra-row spacing was 1 *m*.

Data on VSP plants (2009) were collected during a period of 4 days under well-watered and water-deficit conditions. Water deficit was created by cutting off the irrigation system on the first day of the experiment (July 29^th^). Whole plant transpiration *E_plant_* and net assimilation *A_n, plant_* were monitored using open portable gas-exchange chambers having a cylinder of 1.5 *m* diameter mounted by an open-top cone up to a height of 3 *m*. The frame was made from 4 flat aluminum bars, positioned in a vertical plane on each extremity of the semicircular halves, mounted by 6 semicircular rings in a horizontal plane. The ensemble was covered by a polypropylene film (RXD32 Propafilm, Innovia Films Ltd. UK) of a thickness of 3.5 *μm* and 90% transmission to solar irradiance in the *PAR* band. This plastic material was impermeable to water and CO_2_ and had a negligible water adsorption. Ambient air was taken from above the canopy at 3 *m* height and conducted through flexible plastic pipes covered by a double aluminum foil to avoid solar heating. Air was then injected in the chamber through two holes drilled in the bottom foam and then uniformly distributed through the chamber using holed plastic socks (“plenums”). Air flux was controlled to satisfy a tradeoff between homogenizing air temperature inside the chamber (high flux) and keeping a differential in gas components between the inlets and the outlets (low flux). Air flow entering the chamber was continuously recorded using a differential pressure transducer (PX170, Omega Engineering Inc., UK). Its signal was transformed into flow units through a calibration equation of pressure versus air flow obtained with a reference method.

*E_plant_*, was estimated as:

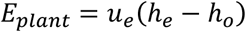

where *E_plant_* is given in [*μmol s*^−1^], *u_e_* is the flow entering the chamber [*mol s*^−1^], and *h_e_* and *h_o_* are respectively the mole fractions of H_2_O of the ambient air at the entry and outlet of the chamber [*μmol mol*^−1^].

Similarly, *A_n, plant_* was calculated as:

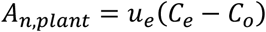

where *A_n, plant_* is given in [*μmol s*^−1^], *C_e_* and *C_o_* are respectively the mole fraction of CO_2_ of the ambient air at the entry and outlet of the chamber [*μmol mol*^−1^].

Temperature of individual leaves inside the chambers were monitored using thermocouples. The latter were inserted into the primary veins of 10 fully-developed individual leaves positioned on different heights from the top of the canopy to the inside, so that temperature gradient resulting from different irradiance conditions may be captured. Temperature was measured continuously and recorded on a 12 minutes basis.

Data on GDC grapevines (2012) were also collected during a 4 days experiment (starting on August 1^st^), but only under water-deficit conditions. Only *E_plant_* rate was monitored by measurements of sap flow using heat energy balance gauges (SGEX25, Dynamax Inc., USA) installed on the two cordons of the GDC plants. The gauges were installed avoiding nodes and irregular trunk sections. Prior to installation, bark was removed to improve the thermal conductivity between the gauges and the sapwood. The gauges were then isolated from air by an aluminum foil which extended down to soil surface in the case of VSP in order to avoid vertical ascending of heat. Heating (5 V) was continuously provided by an external lead-acid battery branched which sent signals to a data logger (CR1000, Campbell Scientific Inc., UK) and a multiplexer (AM16/32, Campbell Scientific Inc., UK). Data was collected every 30 seconds, averaged and recorded every 6 minutes during the experiment.

Stomatal conductance and leaf water potential measurements were performed for a number of leaves on GDC grapevines. *g*_*s, H*_2_*O*_ was measured on three days, at noon, on two fully developed, non-senescent leaves, taken from both sides of the row, using a porometer instrument (AP4, Delta-T Devices, Cambridge, UK). Right after, leaves were cut and introduced in a pressure chamber (Soilmoisture Corp., Santa Barbara, CA, USA) to measure bulk leaf water potential (*Ψ_leaf_*). Measurements of *Ψ_leaf_* were also performed on three days at pre-dawn, in order to estimate soil water status at the beginning of the day, assuming that *Ψ_leaf_* represents *Ψ_soil_* at equilibrium.

### V. Development of the leaf energy balance equation

First, we describe in this section energy balance equations of a planar single leaf whom adaxial (upper) surface is exposed to sky and abaxial (lower) surface is exposed to the soil. Then, we extend the single leaf’s equations to take into account the mutual effects of surrounding leaves on its energy balance.

#### Energy balance of a single leaf

An individual plant leaf gains merely all its energy within both the visible (PAR) shortwave band (400-700 nm) and the infra-red short (NIR) and thermal (TIR) wave bands (respectively 780-3000 nm and 3000-15000 nm). This leaf loses energy by forms of a thermal irradiance within the TIR band, and may further lose energy as a consequence of water evaporation and heat conduction and convection between its surface and surrounding turbulent air masses.

Under the steady state conditions of constant irradiance, air temperature, wind speed and leaf orientation, the gains and losses of energy equilibrate and the energy balance equation of a single leaf at equilibrium writes (**Gutschick, 2016**):

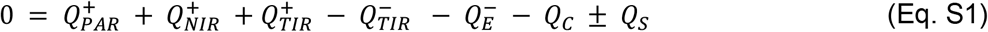

where 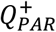 and 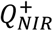 are energy gains respectively in the PAR and NIR wave bands, 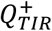 and 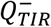 are respectively energy gain and loss in the TIR wave band, 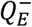 is energy loss accompanying transpiration *E*, *Q_C_* is energy exchange by heat convection and finally *Q_S_* is the energy storage by the leaf. All of the aforementioned energy terms are expressed in units of energy flux density per unit leaf surface area, commonly given in *Wm*^−2^.

Energy input in the shortwave bands PAR and NIR depends on the leaf’s spectral properties: absorptance (*α*), reflectance (*ρ*) and transmittance (*τ*); the three being related (*α* = 1 − *ρ* − *τ*). Energy input writes:

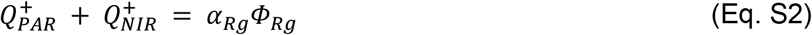

where *α_Rg_* represents the lumped leaf absorptance in the shortwave band. Its value ranges typically from 0.4 to 0.6 **(Nobel, 2005)**.

Energy exchange in the TIR waveband depends weakly on leaf spectral properties, as leaves allows roughly no energy transmittance in this band, while it reflects at most 4% of incident TIR energy fluxes **(Nobel, 2005)**. However, TIR energy exchange becomes heavily dependent on leaf water content which is the dominant TIR-active molecule determining object’s emissivity (*∊*), equal to its absorptivity, in the TIR band. From the Stefan-Boltzmann law for thermal radiation *Q _TIR_* writes:

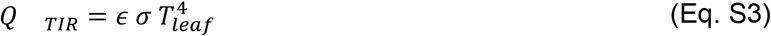

where *σ* is the Stefan-Boltzmann constant [*Wm*^−2^*K*^−4^]. The energy exchange flux densities in the TIR are given by:

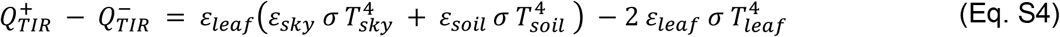

where *ρ_TIR_* is leaf reflectance in the TIR band, *∊_leaf_*, *∊_sky_* and *∊_soil_* are emissivities/absorptivities of the leaf, sky and soil, respectively, and *T_sky_* and *T_soil_* are respectively the absolute temperatures of the sky and soil [*K*]. In the above equation, TIR energy loss occurs from both surfaces of the leaf 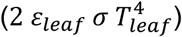 while it is assumed that sky input 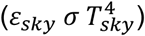 and soil input 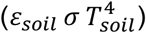 are attributed to adaxial and abaxial surfaces, respectively, in the absence of transmittance.

Regarding the energy loss due to water transpiration, this term writes:

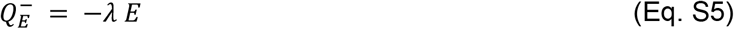

where *λ* is latent heat for vaporization [*W s mol*^−1^].

The energy exchange by conduction-convection can be deduced from Fick’s first law:

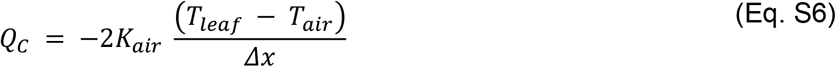

where *K_air_* is the thermal conductivity coefficient of air [*W m*^−1^*K*^−1^].

Finally, the energy storage term is *Q_S_* englobes the energy used by the metabolic reactions within the leaf and the energy stored because of changes in leaf temperature. This term is negligible since both phenomena produce energy changes that are account for less than 1% of the sole gain of energy in the shortwave band **(Nobel, 2005)**. It is hence assumed in this work that *Q_S_* equals 0.

Combining equations S1 to S6 yields the following general equation for energy balance of a single leaf:

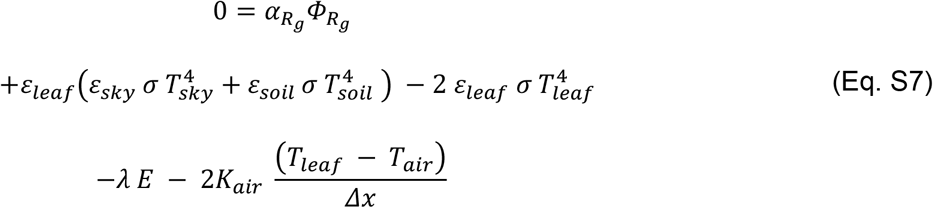

#### Energy balance of a leaf inside a canopy

Leaf positioning inside the canopy affects its energy input, both in the visible and infrared wave bands. Energy exchange between leaves can be accounted for by the radiosity method, or using one of its derivatives as for the nested radiosity employed by the Caribu light interception model **(Chelle et al., 1998)**.

The incident energy on an individual leaf within a canopy comes from direct sources (sun, sky and clouds) and from energy that has been scattered by the surrounding elements (typically plant shoots and soil). Both direct and scattered energy are weighted by their solid angle that subtends the the leaf in question:

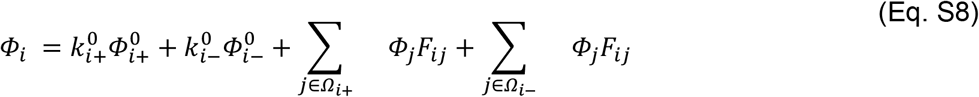

where *Φ_i_* is the incident energy reaching the leaf *i*, 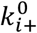 and 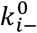 are form factors proportions of the solid angles of direct energy sources reaching respectively the adaxial (denoted *i* +) and abaxial (*i* −) surfaces, 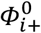 and 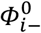 are energy flux densities reaching *i* + and *i* −, respectively, *Ω*_*i*+_ and *Ω*_*i*−_ are two hemispheres covering *i* + and *i* −, respectively, *E_j_* is the scattered energy coming from a leaf *j*, and finally *F_ij_* is the form factor defined as:

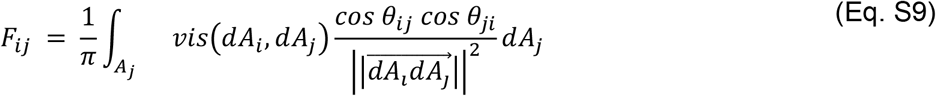

where *θ_ij_* the angle between the normal to the leaf *i*’s surface, *A_i_*, and the the direction 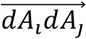, *θ_ij_* is the angle between the normal to leaf *j*’s surface *A_j_* and the direction 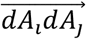, *vis*() is a binary function whose output is 1 when no occlusion occurs between *dA_i_* and *dA_j_*, otherwise it yields 0.

Equations S8 and S9 apply to energy fluxes in both visible and infrared bands. Applying the appropriate changes to Eq.S7 yields the general equation for energy balance of an individual leaf inside a canopy:

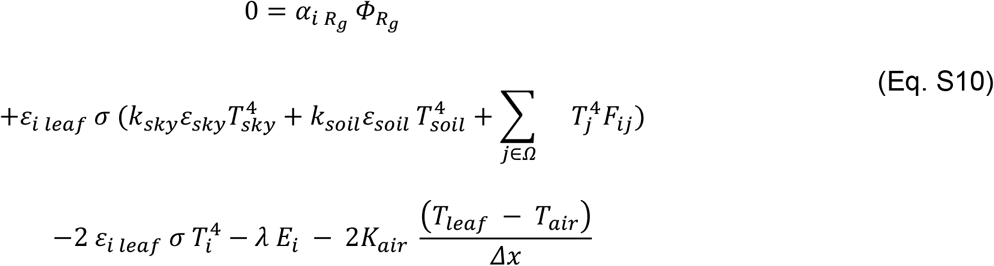

where the indice *i* has been added to distinguish the parameters specific to the leaf in question.

